# LRRC37B is a species-specific regulator of voltage-gated channels and excitability in human cortical neurons

**DOI:** 10.1101/2022.12.21.521423

**Authors:** Baptiste Libé-Philippot, Amélie Lejeune, Keimpe Wierda, Ine Vlaeminck, Sofie Beckers, Vaiva Gaspariunaite, Angéline Bilheu, Hajnalka Nyitrai, Kristel M. Vennekens, Thomas W. Bird, Daniela Soto, Megan Y Dennis, Davide Comoletti, Tom Theys, Joris de Wit, Pierre Vanderhaeghen

## Abstract

The enhanced cognitive abilities characterizing the human species result from specialized features of neurons and circuits, but the underlying molecular mechanisms remain largely unknown. Here we report that the hominid-specific gene *LRRC37B* encodes a novel receptor expressed in a subset of human cortical pyramidal neurons (CPNs). LRRC37B protein localizes at the axon initial segment (AIS), the specialized domain triggering action potentials. *LRRC37B* ectopic expression in mouse CPNs *in vivo* leads to reduced intrinsic excitability, a distinctive feature of some classes of human CPNs. At the molecular level, LRRC37B acts as a receptor for the secreted ligand FGF13A and interacts with the voltage gated sodium channel (VGSC) beta subunit SCN1B, thereby inhibiting the channel function of VGSC, specifically at the AIS. Electrophysiological recordings in adult human cortical slices reveals that endogenous expression of LRRC37B in human CPNs reduces neuronal excitability. *LRRC37B* thus acts as a species-specific modifier of human cortical neuron function, with important implications for human brain evolution and diseases.

## Introduction

Humans display distinctive cognitive abilities compared to other animals including our closest living hominid relatives, chimpanzees and bonobos (Richerson et al., 2021). These specialized functions are thought to rely on features that distinguish the human brain from other species, in particular at the level of the cerebral cortex. Distinctive properties of the human cortex include an increased number of cortical neurons, an increased proportion of specific neuronal classes (such as cortical pyramidal neurons, (CPNs) populating the upper layers and the prefrontal areas), as well as increased numbers of synapses between cortical neurons, or cortico-cortical connectivity (Buckner and Krienen, 2013; Herculano-Houzel, 2009; Schmidt and Polleux, 2022; Sousa et al., 2017). These changes have been linked to the evolution of developmental mechanisms controlling the generation and specification of cortical neurons and circuits, which start to be unravelled by convergent work of human genomics, genetics and embryology, as well as pluripotent stem cell-based modelling (reviewed in (Bae et al., 2015; Libé-Philippot and Vanderhaeghen, 2021; Sousa et al., 2017).

In addition to these global changes, the evolution of human brain circuits may also be linked to divergent features at the level of individual neurons, as described in other species (Roberts et al., 2022). Human CPNs have long been known to display distinctive morphological properties, including larger and more branched dendritic arbours (Sherwood et al., 2003) and increased numbers of synapses per neuron compared to all other primates (Sherwood et al., 2020). Moreover, recent work using electrophysiological recordings of live *ex vivo* human and non-human brain sections has uncovered functional properties of human CPNs that distinguish them from other species. These include specialized synaptic features leading to stronger synapses (Molnár et al., 2008, 2016), displaying higher levels of plasticity (Szegedi et al., 2016) or recovery (Testa-Silva et al., 2014). Some classes of human CPNs also display distinct biophysical and intrinsic properties, which have been linked to the unusually large size of their dendritic arbors, but also differences in membrane properties, including distinct distribution and density of ion channels (Beaulieu-Laroche et al., 2018, 2021; Deitcher et al., 2017; Eyal et al., 2016; Geirsdottir et al., 2019; Gidon et al., 2020; Kalmbach et al., 2018, 2021; Schwarz et al., 2019).

These electrophysiological differences characterizing similar neuronal subtypes between species have been proposed to have significant consequences on neuronal function (Beaulieu-Laroche et al., 2018, 2021; Deitcher et al., 2017; Eyal et al., 2016; Geirsdottir et al., 2019; Gidon et al., 2020; Kalmbach et al., 2018, 2021; Schwarz et al., 2019) and circuit processing (Itoh et al., 2022; Woodman et al., 2007). Interestingly, they could account for specialization of neuronal intrinsic excitability, which is indeed decreased in some classes of human CPNs compared with other species (Beaulieu-Laroche et al., 2018, 2021). Excitability is a critical regulator of neuronal and circuit function, as it determines how neurons integrate external synaptic influences, or input, to generate specific action potential trains at the axonal level, or output. Input/output relationship can critically affect information processing and neural circuit plasticity, for instance through neuronal gain modulation, by which neurons adapt to changing inputs, through Hebbian or homeostatic mechanisms (Beck and Yaari, 2008; Ferguson and Cardin, 2020).

The recently uncovered changes in neuronal intrinsic properties could thus contribute in an important fashion to the enhanced memory and information processing that characterize our species, but the underlying cellular and molecular mechanisms remain essentially unknown.

At the cellular level, the input/output function is known to be modulated at the level of all neuronal compartments, from dendrites to soma and axon, and at the level of the axon initial segment (AIS), the specialized axonal compartment site of action potential generation (Debanne et al., 2019). In particular, the size and location of the AIS, the density and modulation of ion channels concentrated at its level, such as voltage-gated sodium channels (VGSC), are critical components of neuronal excitability (Grubb and Burrone, 2010; Kole et al., 2008; Leterrier, 2016). Intriguingly, while the changes of human CPNs have been so far essentially linked to synaptic and dendritic features, whether and how the AIS contributes to human neuron specialization remains unknown.

At the molecular level, the specialization of human cortical neurons could be driven by evolutionary changes in the gene regulatory networks underlying their identity (Lein et al., 2017; Tosches, 2021). Evolutionary novelties in human neurons could also emerge from new genes arising through recent gene duplication in the human lineage (Dennis and Eichler, 2016). Some of these hominid-specific (HS) gene duplicates were previously identified as human-specific modifiers of human brain development (Libé-Philippot and Vanderhaeghen, 2021).

Here we focus on *LRRC37B*, a member of the HS family of Leucine Rich Repeat (LRR) orphan receptors *LRRC37* (Giannuzzi et al., 2012). We find that LRRC37B protein is uniquely displayed at the level of the AIS in human CPNs. LRRC37B negatively regulates intrinsic excitability following overexpression in the mouse, and endogenous expression of LRRC37B in human CPNs controls neuronal excitability. Molecularly, LRRC37B acts as a co-receptor for FGF13A and SCN1B, and thereby inhibits the function of voltage-gated sodium channels, specifically at the AIS level. Our data identify a hominid-specific receptor and its interactors that regulate human neuronal properties in a human-specific fashion, thereby shedding light on the molecular mechanisms of human neuronal evolution.

## Results

### *LRRC37* gene family is selectively expressed in human cortical neurons

We previously identified >30 HS genes expressed during human corticogenesis (Suzuki et al., 2018), among which the *LRRC37* (Leucine-Rich Repeat-containing protein 37) gene family that encodes putative membrane receptors (Giannuzzi et al., 2012). LRRC37 receptors have no known function or ligands, but interestingly they belong to the larger structural group of Leucine-Rich Repeat (LRR) receptors, previously implicated in key aspects of neuronal development and connectivity (De Wit et al., 2011).

The number of *LRRC37* genes increased dramatically in simian species leading to many paralogs (Giannuzzi et al., 2012). Among them, the chimpanzee and human genomes carry at least 4 protein-encoding genes, compared to 2 in the mouse genome (Figure 1A) (Giannuzzi et al., 2012). *LRRC37* genes encode two types of proteins, A and B. Both A-and B-types are transmembrane proteins contain a Leucine-Rich Repeat (LRR) domain in the extracellular core, but the B-type carries an additional specific domain, hereafter referred to as LB domain, located near the N-terminal part of the protein (Figure 1B). *LRRC37B* emerged in the simian genomes (Figure 1A). Alignment of the amino acid sequences of *LRRC37B-type* genes found in primates revealed strong conservation between human and chimpanzee, especially at the level of the LB and LRR domains (96-100%). On the other hand the macaque *LRRC37B*-type gene (LRRC37-M2) only displays lower levels (63-80%) of homology, and its LRR domain contains only three LRR, instead of six in human and chimpanzee (Figure 1, Figure S1A). The *LRRC37B* gene studied here thus appears to be present only in human and chimpanzee genomes. Finally and importantly, when examining copy-numbers (CN) of *LRRC37* human paralogs predicted to encode proteins using whole-genome sequencing read depth (Shen and Kidd, 2020), *LRRC37A* and *LRRC37A2* exhibit polymorphism (diploid CN between 0 and 5) while in contrast *LRRC37A3* and *LRRC37B* are near fixed (diploid CN = 2) across thousands of diverse humans from the 1000 Genomes Project (Auton et al., 2015) (Figure S1B).

**Figure 1.**
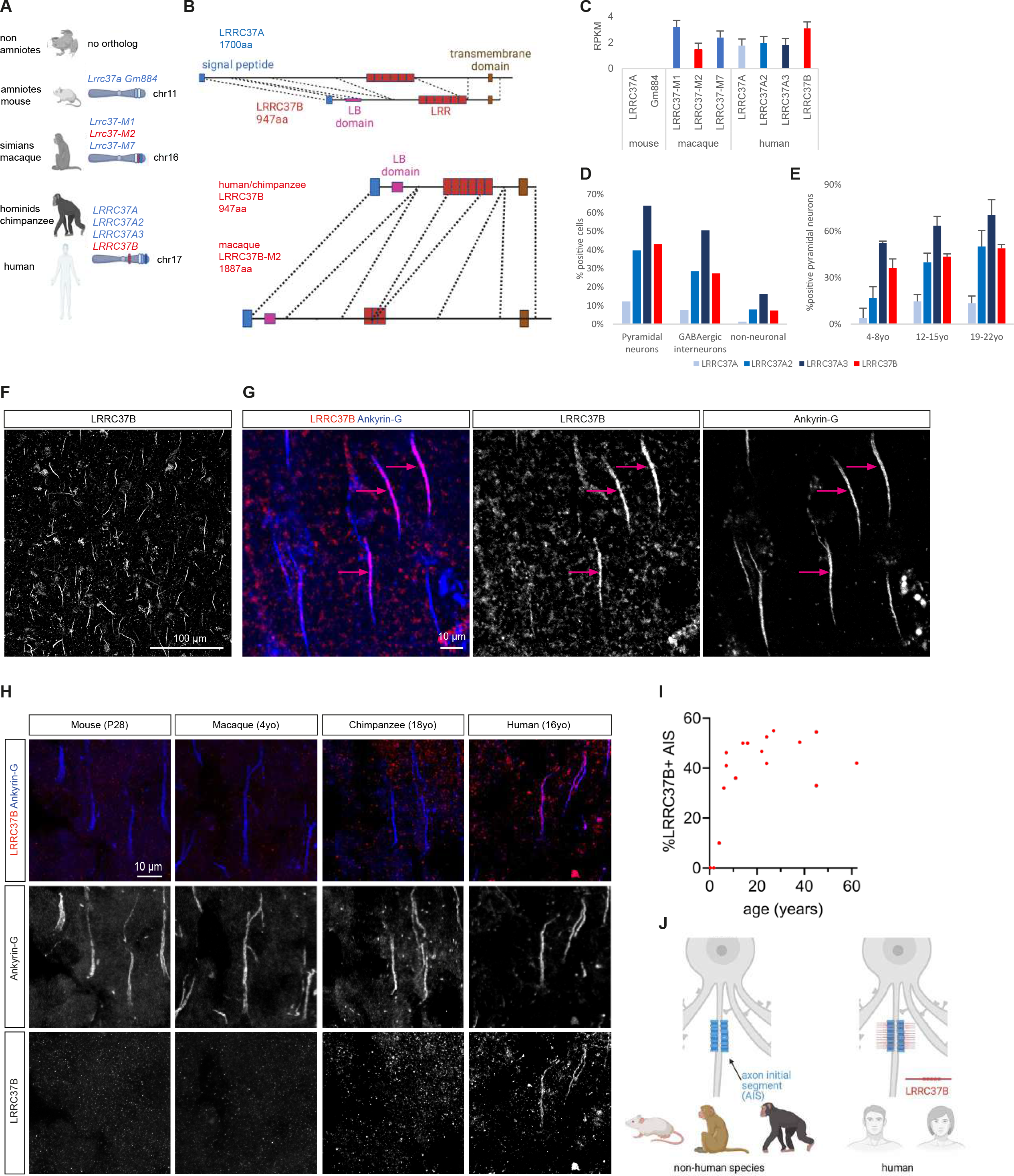
LRRC37 gene family evolution and expression in mammalian cortex. **A,** Orthologs and paralogs of the *LRRC37* genes in mouse, macaque and hominids (from (Giannuzzi et al., 2012) and Ensembl); A-types in blue, B-types in red. **B,** Protein structure of human LRRC37 proteins as well as other simian LRRC37B proteins; note the presence of a LRRC37B specific “LB domain”. **C,** *LRRC37* transcripts detected by bulk RNAseq in the cerebral cortex in mouse, macaque and human (from(Cardoso-Moreira et al., 2019), RPKM: Reads Per Kilobase Million; median + SE median). **D-E**, Proportion of cell types and pyramidal neurons (CPN) at different ages (years old, yo) with *LRRC37* transcripts detection from single cell RNAseq from human postnatal cortical samples (yo: years old, from (Velmeshev et al., 2019), mean + SEM); non-neuronal cells include astrocytes, microglia, endothelial cells and oligodendrocytes. **F-G**, Immunofluorescent detection of LRRC37B in the human cerebral cortex using an antibody directed to the LB domain: LRRC37B colocalizes with a subset of axon initial segments (AIS) (G, arrows: LRRC37B+ neurons) of CPN labelled by Ankyrin-G. **H**, Immunofluorescent detection of LRRC37B and Ankyrin-G in the mouse (postnatal day 28, P28) barrel cortex, macaque, chimpanzee and human temporal cerebral cortex (cryosections): LRRC37B is uniquely detected in the human cerebral cortex. **I**, Proportion of CPNs positive for LRRC37B at the level of the AIS from neonates to 61yo: LRRC37B is detected in the human cerebral cortex from 2yo to reach 35-60% of positive AIS between 10yo and 61yo. **J**, Schematic summary: LRRC37B is a protein located at the AIS specifically in human.

Inspection of bulk RNAseq datasets (Cardoso-Moreira et al., 2019) and quantifying expression using an approach sensitive to duplication genes (Suzuki et al., 2018) revealed that all *LRRC37* paralogs are robustly expressed in the human and macaque developing and adult cerebral cortex, while no *LRRC37* transcripts are detected in the mouse cortex (Figure 1C). Human *LRRC37B* is expressed at higher levels at the RNA level than the *LRRC37B-type* gene in macaque cortex (Figure 1C) in line with previous findings (Giannuzzi et al., 2012). Inspection of scRNAseq data (Velmeshev et al., 2019) revealed that *LRRC37* genes are expressed in most classes of CPNs and interneurons, but only in a subset of neurons in each subclass (Figure 1D, Figure S1C-D). The proportion of LRRC37B-positive CPNs is very low at birth, and then gradually increases during the first two decades of life to reach a plateau, variable for each subclass, in the adult human cortex (Figure 1E, Figure S1D).

### LRRC37B is displayed at the axon initial segment of human cortical pyramidal neurons

We next focused on LRRC37B, as it constitutes the most recent genomic event in the hominid lineage, and its copy number appears to be nearly fixed in the human population, suggesting conservation and functionality (Figure 1A, Figure S1B). We first examined LRRC37B at the protein level, using a specific antibody raised against the LB domain. This antibody recognizes LRRC37B-type proteins encoded in human, chimpanzee and macaque, but not the LRRC37A-type (Figure S1E-F).

Using this antibody, we confirmed that LRRC37B can be targeted to the cellular membrane with the expected topology in heterologously transfected HEK-293T cells, using immunostaining before permeabilization (Figure S1E). We next examined the pattern of endogenous LRRC37B expression in the adult human cerebral cortex. Remarkably, this revealed that LRRC37B is mostly localized at the level of the axon initial segment (AIS, marked with ankyrin-G) in a subset (>50%) of pyramidal neurons (Figure 1F-I, Table 1). The AIS subcellular compartment is the main site of generation of action potentials (AP) through the opening of VGSC alpha subunits (NAVα) concentrated at this location (Leterrier, 2016). Interestingly, similar immunostaining on adult cerebral cortex of mouse, macaque and chimpanzee with the same antibody failed to reveal any detectable expression (Figure 1H). Time-course analysis at several stages from early postnatal ages to >60 years old human specimens, revealed that the proportion of neurons displaying LRRC37B at the AIS sharply increased from birth until childhood to stabilize at puberty (Figure 1I, Table 1), in good concordance with the RNA levels (Figure 1E).

Overall these data identify LRRC37B as a novel LRR membrane-anchored protein selectively expressed at the AIS of human cortical pyramidal neurons (Figure 1J).

### LRRC37B negatively regulates cortical pyramidal neuron excitability

In order to probe the function of LRRC37B, we next performed sparse gain-of-function in mouse pyramidal neurons through sparse in utero electroporation in the cortex (at embryonic day 15.5, thereby targeting mostly cortical layer 2-3 pyramidal neurons). We thereby electroporated vectors leading to a cre-dependent bicistronic expression of LRRC37B and EGFP, or EGFP alone as a control, followed by analysis at P28 (see Methods). We found that the LRRC37B protein was enriched at the AIS, as in human neurons (Figure 2A, Figure S2A), while control neurons displayed no LRRC37B immunoreactivity, as expected (Figure S2A). Notably we detected no differences in the AIS length and localization in LRRC37B-expressing neurons, compared with control neurons (Figure S2B-C). Interestingly, ectopic expression of the more ancestral gene LRRC37A2 did not reveal a selective localization at the AIS (Figure S2D), further pointing to novel properties of the *LRRC37B* paralog.

**Figure 2.**
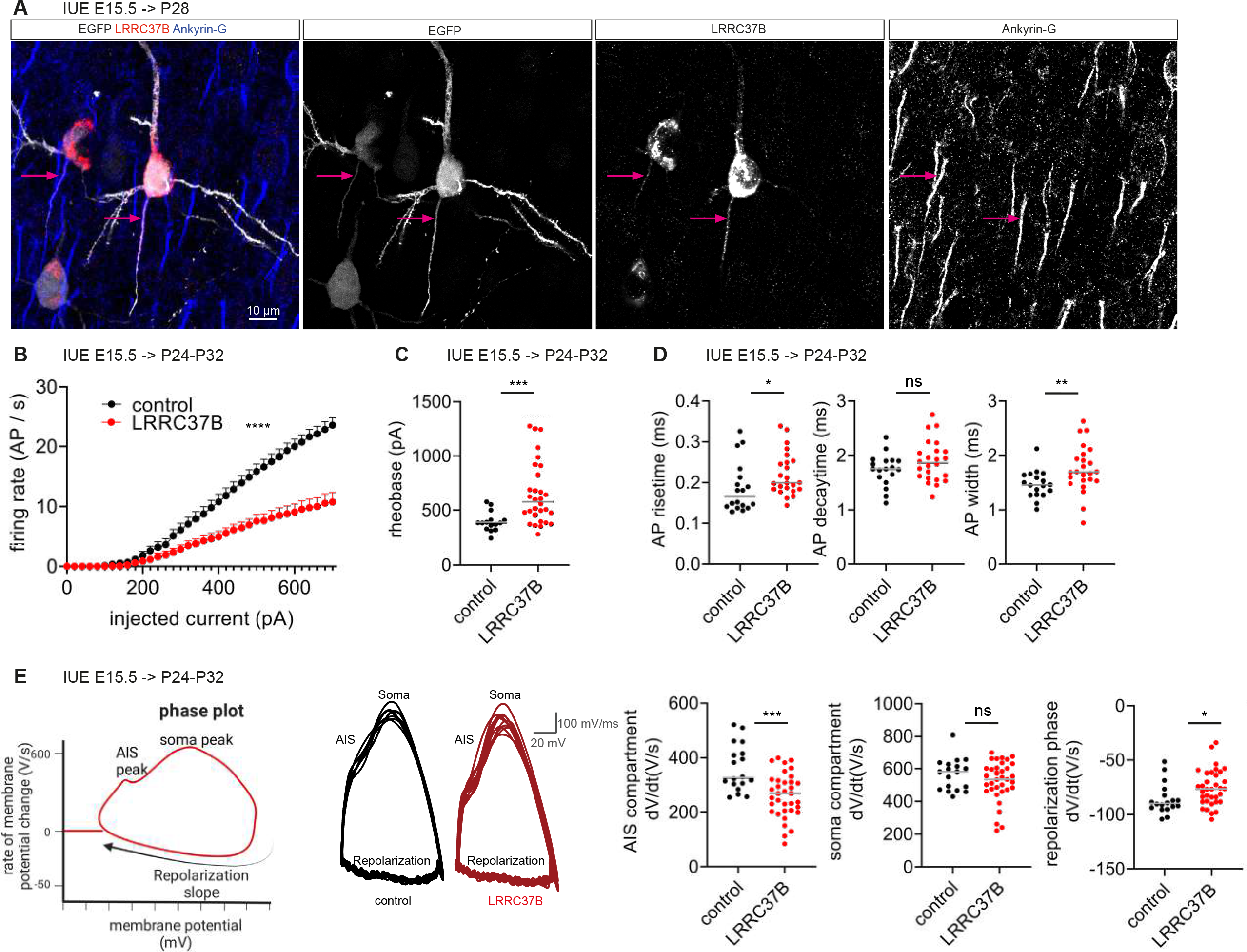
LRRC37B is a human AIS protein sufficient to induce a decreased neuronal excitability. **A,** Immunodetection of LRRC37B (arrows) and Ankyrin-G in the mouse cerebral cortex after transfection of LRRC37B and EGFP cDNAs. **B-C,** Intrinsic excitability of pyramidal neurons transfected for LRRC37B and EGFP cDNAs compared to EGFP cDNA only (2-way ANOVA test for action potential (AP) firing rate, Mann-Whitney test for rheobase). **D**, Action potential (AP) properties of pyramidal neurons transfected for LRRC37B and EGFP cDNAs compared to EGFP cDNA only (Mann-Whitney test). **E**, Phase plot analysis of single APs (Mann-Whitney tests). ns, non-significative; *; p<0.05; **; p<0.01; ***, p<0.001; ****, p<0.0001.

Given the prominent role of the AIS in controlling neuronal excitability, we next compared the electrophysiological properties of control and LRRC37B-expressing mouse cortical CPNs, using patch-clamp recordings in *ex vivo* cortical slices of electroporated mice at P24-P32. Remarkably, *LRRC37B* gain-of-function led to a sharp decrease in neuronal excitability, characterized by a lower AP firing rate and a higher rheobase (Figure 2B-C, Figure S2E), while sodium currents remained unchanged (Figure S2F). It also led to a decreased input resistance and an increased capacitance (Figure S2E), which could in principle account for changes in the firing rates. However, LRRC37B-expressing neurons also displayed specific changes in two critical features of AP, risetime and width, pointing to changes in intrinsic excitability of the neurons (Figure 2D). This was confirmed by phase plot analyses of single AP, which revealed that LRRC37B-expressing neurons display a decrease in the AIS compartment of the AP generation (-27.7%), while the somatic compartment remained largely unchanged (Figure 2E). Phase plots also revealed a decrease in the repolarization phase that occurs after the peak of the AP in LRRC37B-expressing neurons (Figure 2E).

We next examined the synaptic properties of LRRC37B-expressing neurons using whole-cell voltage clamp recordings, which revealed no obvious difference in frequency or amplitude of spontaneous excitatory and inhibitory postsynaptic currents compared with control-electroporated neurons (Figure S2G). As the AIS is the synaptic target of chandelier GABAergic interneurons that thereby regulate CPN output (Gallo et al., 2020), we quantified GABAergic synapses at the level of the AIS. This revealed no difference between LRRC37B-expressing and control mouse CPNs in the density of vGAT or Gephyrin puncta at the level of the AIS (Figure S2H-I).

These data indicate that LRRC37B can act as a negative regulator of intrinsic excitability when overexpressed in mouse CPNs, acting mostly at the level of AP generation at the AIS.

### LRRC37B is a membrane receptor for extracellular FGF13A, which binds to voltage-gated sodium channels at the AIS

In order to identify the molecular mechanisms underlying the function of LRRC37B in neuronal excitability, we searched for LRRC37B binding partners. We first performed an ELISA-based unbiased interactome screen (Ozgul et al., 2019; Ranaivoson et al., 2019). For this approach, we used as a bait the predicted extracellular sequence of LRRC37B (LRRC37B_ECTO) fused with alkaline phosphatase (AP) (Apóstolo et al., 2020). Using LRRC37B_ECTO-AP immobilized in each well of 384-well plates, 920 transmembrane or secreted proteins (including 23 proteins enriched at the AIS or in chandelier interneurons, see Methods), fused to an Fc domain, were used as preys. The screen led to a single reproducible hit from three independent experiments: FGF13 isoform 1 (hereafter named FGF13A) (uniprot ID: Q92913-1). This result came first as a surprise, as FGF13 was previously described as a member of the FHF non-canonical FGF family, thought to encode non-secreted proteins (Dover et al., 2010; Smallwood et al., 1996; Wang et al., 2012). Interestingly, FGF13 encodes several splice isoforms (FGF13A, FGF13B, FGF13V, FGF13Y and FGF13VY) (Figure 3A) that share a C-terminal domain (encoded by coding exons 2-5, thereafter named “core domain”) but differ by their first coding exon (Munoz-Sanjuan et al., 2000).

**Figure 3.**
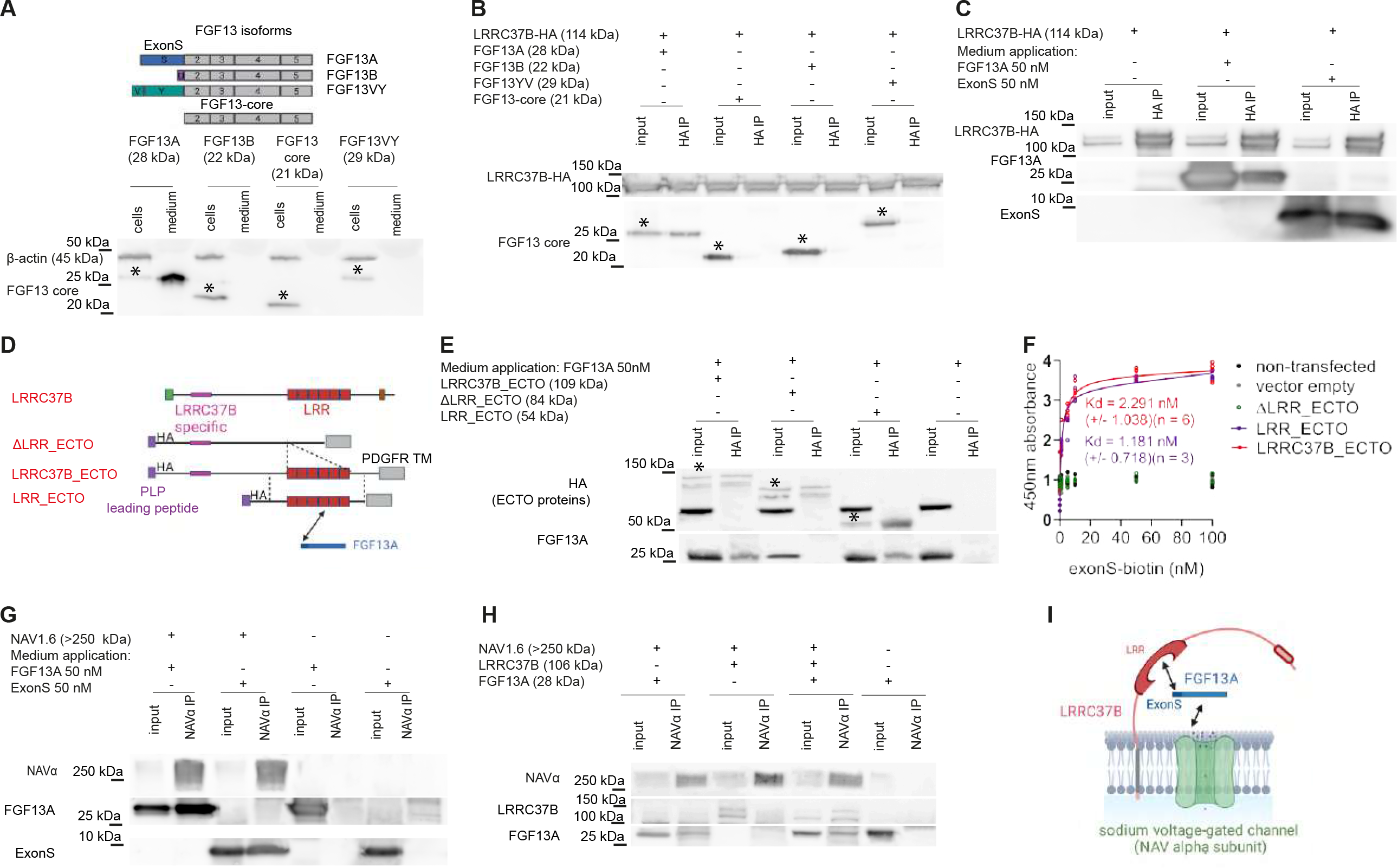
LRRC37B is a receptor for FGF13A. **A**, FGF13 codes for several spliced isoforms; cell lysate and medium samples of HEK-293T cells transfected for FGF13A, FGF13B, FGF13VY and FGF13-core cDNAs (stars indicate in the cell extracts for each isoform). **B**, LRRC37B-HA immunoprecipitation from HEK-293T cells co-transfected for LRRC37B-HA cDNA and cDNAs coding for the different FGF13 isoforms (stars indicate in the input for each isoform). **C**, LRRC37B-HA immunoprecipitation from HEK-293T cells transfected for LRRC37B-HA cDNA and with recombinant FGF13A or its synthetic ExonS applied in the culture medium. **D-E**, Immunoprecipitation of different protein lacking or carrying the leucin-rich repeats (LRR) of the extracellular domain of LRRC37B from transfected HEK-293T cells with recombinant FGF13A application in the culture medium (stars indicate in the input each LRRC37B protein). **F,** Binding assay of synthetic FGF13A ExonS-biotin to the different LRRC37B proteins carrying or lacking LRR expressed by transfected HEK-293T cells (fitting curves for LRRC37B_ECTO and LRR_ECTO, see Methods). **G**, Immunoprecipitation of NAV1.6 (SCN8A) from HEK-293T cells transfected for SCN8A/NAV1.6 cDNA with recombinant FGF13A or its synthetic ExonS applied in the culture medium. **H**, NAV1.6 immunoprecipitations from HEK-293T cells transfected for SCN8A/NAV1.6 +/-FGF13A +/- LRRC37B cDNAs. **I**, Schematic summary of the LRRC37B – FGF13A – NAV1.6 interaction.

We hypothesized that alternatively spliced mRNAs could encode secreted or non-secreted isoforms of FGF13. We tested this using heterologous expression in HEK cells of all major FGF13 isoforms, followed by analysis of cell lysates vs. medium. This revealed that only FGF13A can be detected as a secreted protein in the culture medium, but none of the other tested isoforms, nor the core domain (Figure 3A).

We next tested whether FGF13A could bind to LRRC37B in the same system. LRRC37B was found to co-immunoprecipitate only with FGF13A, and not with any of the other tested isoforms, nor with FGF13 core domain (Figure 3B). This interaction was further confirmed by adding recombinant FGF13A protein to the culture medium of cells expressing *LRRC37B*, which also resulted in co-immunoprecipitation of FGF13A with LRRC37B (Figure 3C). Finally, and importantly, similar binding results were obtained with a synthetic peptide corresponding to the first exon of FGF13A (ExonS, Figure 3C), indicating that the ExonS-encoded domain alone is sufficient to bind to LRRC37B.

Collectively these data indicate that FGF13A is an extracellular ligand of LRRC37B, to which it binds to through its isoform-specific N-terminal domain.

In order to identify the domains of LRRC37B receptor responsible for the interaction with FGF13A, we generated a series of LRRC37B constructs containing deletions in the extracellular domain of LRRC37B, fused at the N-terminal with the prolactin leader peptide and an HA tag, and at the C-terminal with the transmembrane domain of PDGF-R (Figure 3D). We performed immunoprecipitations of the different LRRC37B deletion mutants transfected in HEK-293T cells, following application of FGF13A to the culture medium. This revealed that the LRR domain (amino acids 468 – 841) is necessary and sufficient to bind to FGF13A (Figure 3E). Similar co-immunoprecipitations were observed following co-transfection of LRRC37B and FGF13A cDNAs (Figure S3A). We also compared the molecular properties of human LRRC37B with macaque and chimpanzee LRRC37B proteins. This revealed that the chimpanzee protein could bind to FGF13A, while the macaque LRRC37B did not, in line with its shorter LRR domain structure (Figure1B, Figure S3C). Similarly, human LRRC37A2, which displays an LRR domain very similar to LRRC37B (Figure S1A), could bind to FGF13A (Figure S3B). To assess the specificity of the interaction between LRRC37 proteins and FGF13A, we performed the same approach with the extracellular domains of other membrane receptors, including several LRR receptors (GPR158, SLITRK2, LRRTM1 and CD4). None of them co-immunoprecipitated FGF13A (Figure S3C).

Collectively these data indicate that LRRC37B binds to FGF13A through its LRR domain. To estimate the affinity of the interaction between FGF13A and LRRC37B, we performed binding assays using a biotin-labelled FGF13A-ExonS-peptide and HEK-293T cells expressing LRRC37B and deletion mutants (Figure 3F). This revealed an apparent high affinity specific binding of FGF13A-ExonS-peptide to the extracellular domain of LRRC37B (Kd:2.291+/- 1.038 nM) and to the LRR domain alone (Kd:1.181 +/- 0.718nM), while no binding was detected with LRRC37B mutant devoid of its LRR (Figure 3F).

FGF13 was previously reported to modulate the function of VGSC NAVα subunits, but through binding of its core domain to an intracellular domain of the channel (Barbosa et al., 2017; Goetz et al., 2009; Wang et al., 2011; Wittmack et al., 2004). Given our data above, we hypothesized that FGF13A could potentially bind to NAV channels extracellularly. We first tested whether FGF13A and its ExonS could bind to NAV channel alpha subunit NAV1.6, a major NAV alpha subunit that initiates AP at the AIS of pyramidal neurons (Hu et al., 2009; Leterrier, 2016; Lorincz and Nusser, 2008). Strikingly, we found that FGF13A, or its ExonS only, when applied in the medium of HEK cells transfected for NAV1.6, could be immunoprecipitated with NAV1.6 (Figure 3G). We next tested the relevance of these observations to LRRC37B by co-transfecting the three proteins in HEK-293T cells. Importantly, this revealed that LRRC37B could not be co-immunoprecipitated with NAV1.6 alone, while it could be found in the same complex in the presence of FGF13A (Figure 3H).

Our data thus lead to a molecular model whereby extracellular FGF13A binds directly to the NAV channel alpha subunit NAV1.6 and to LRRC37B, and the three proteins can be found in the same complex when co-expressed together (Figure 3I).

### Extracellular FGF13A decreases cortical neuron excitability and is concentrated at the AIS by LRRC37B

We then explored the relevance of our molecular findings in the mouse cortex *in vivo*. Notably, FGF13A, including its ExonS peptide, and NAV1.6 amino acid sequences are highly conserved between mouse and human species (Table 2). We first performed co-immunoprecipitation of LRRC37B following mouse cerebral cortex IUE as above, which confirmed the interaction *in vivo* between LRRC37B and FGF13A (Figure 4A). We next performed immunostainings for LRRC37B, FGF13A and NAV1.6 in mouse cortical sections at P28, following in utero-electroporation of LRRC37B and eGFP or control eGFP, and analysed these using STED microscopy. This revealed that LRRC37B co-localizes with NAV1.6 and FGF13A at the AIS (Figure 4B). Furthermore, following LRRC37B transfection, FGF13A was found to be significantly more abundant at the AIS, in distinct patches, co-localized with LRRC37B (Figure 4C-D).

**Figure 4.**
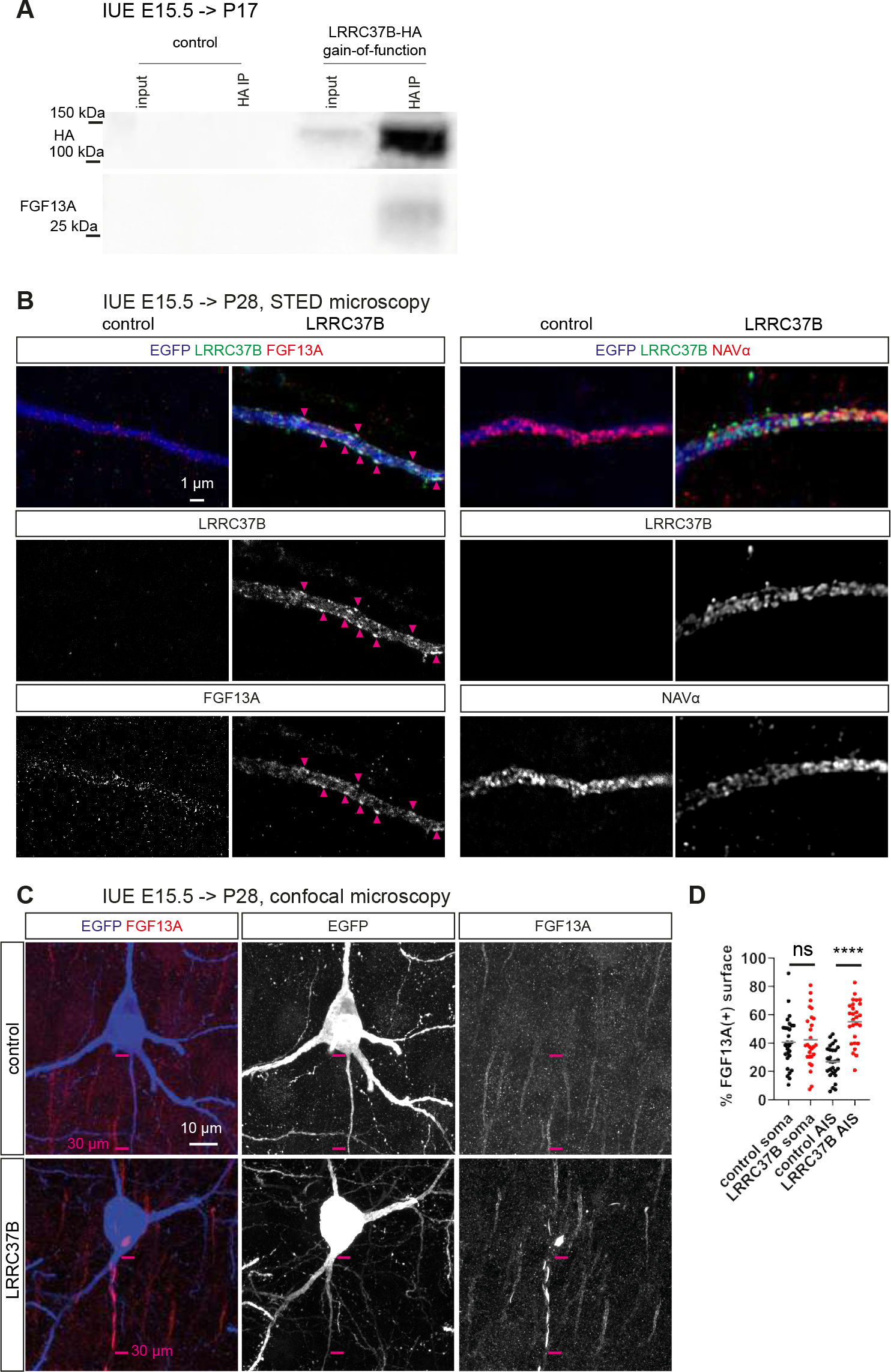
LRRC37B interacts with FGF13A *in vivo* and enhanced its amount at the AIS. **A**, Immunoprecipitation of LRRC37B-HA from mouse cortical protein extract (P17) of LRRC37B-HA/EGFP transfected mouse cortex compared to EGFP alone. **B**, STED microscopy of the AIS of mouse neurons transfected for LRRC37B/EGFP or EGFP alone immunostained for EGFP, LRRC37B, FGF13A and NAVα subunits; arrowheads indicate colocalizations between LRRC37B & FGF13A. **C,** Mouse neurons transfected for LRRC37B/EGFP or EGFP alone immunostained for EGFP and FGF13A. **D**, Corresponding quantification of FGF13A staining at the soma and AIS levels (Mann-Whitney tests; ns, non-significative; ****, p<0.0001).

Together, these observations indicate that LRRC37B colocalizes with FGF13A and NAV1.6 at the AIS, and concentrates FGF13A at the AIS in vivo.

We next tested whether and how FGF13A may affect the physiology of the neurons. To this aim we first performed patch-clamp recordings of mouse non-electroporated cortical slices (at P24-P32) combined with the extracellular bath application of FGF13A (Figure 5A-D, Figure S4A-E). This revealed a dose-dependent effect of FGF13A on neuronal excitability at 10-50 nM, leading to decreased AP firing rate, increased rheobase, increased AP risetime and width, thus strikingly mimicking the effects of LRRC37B gain of function (Figure 5A-C, Figures S4A-D). Notably, FGF13A addition had no effect on membrane resistance and capacitance. Phase plot analyses of single AP revealed that FGF13A extracellular application led to a decrease in the AIS compartment (-17.8%) and repolarization phase, as observed following LRRC37B gain of function. Interestingly, it also led to a decrease in the soma compartment (-9.4%) (Figure 5D) and to a global decrease in sodium currents (Figure S4E), two effects not seen following LRRC37B overexpression (Figure 2E, Figure S2E).

**Figure 5.**
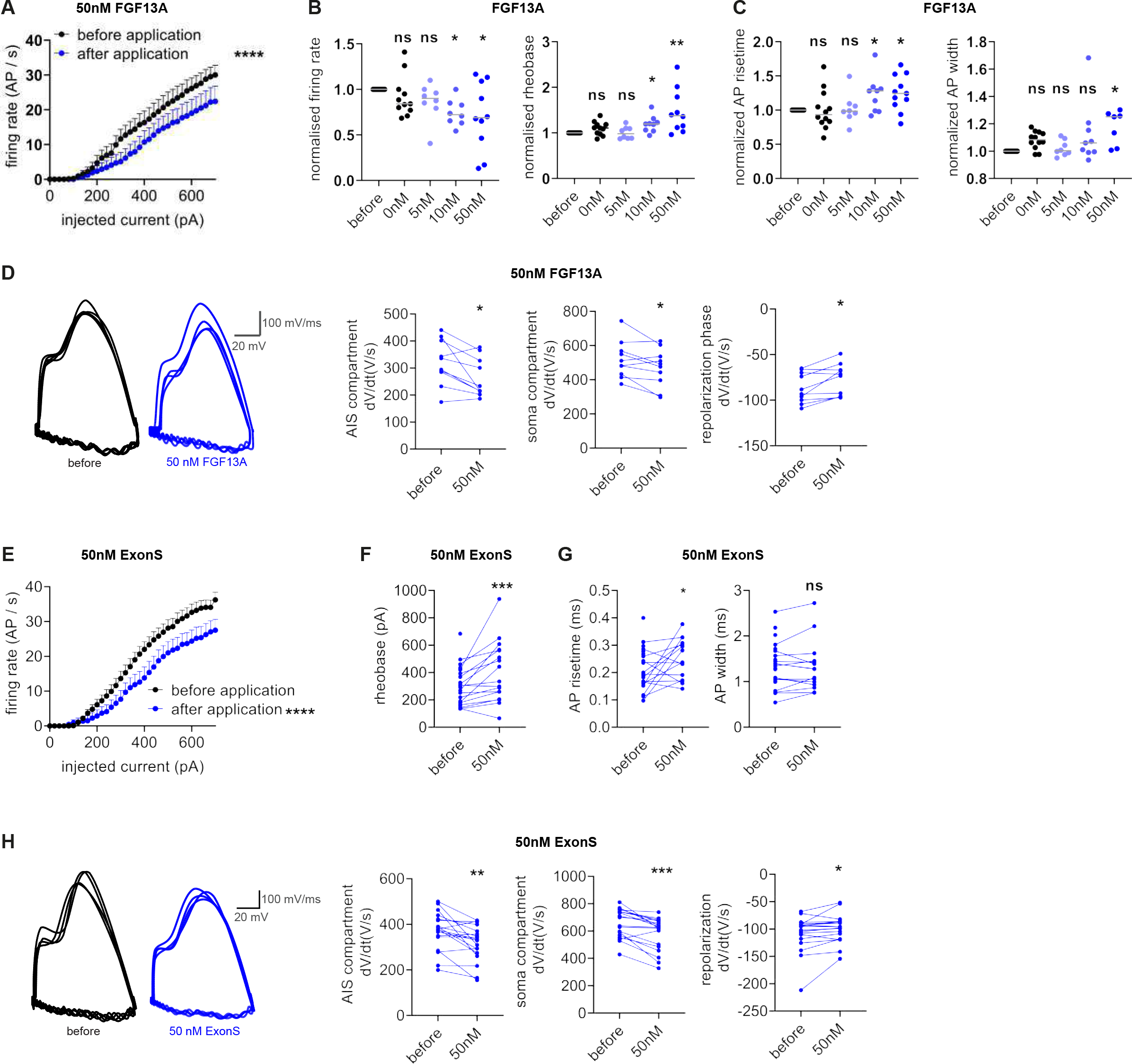
FGF13A regulates neuronal excitability through its ExonS. **A**, AP firing rate of mouse pyramidal neurons before and after 50nM recombinant FGF13A extracellular application (2-way ANOVA test). **B-C**, Dose-response effect of recombinant FGF13A extracellular application on cell intrinsic properties (AP firing rate normalized at 700 pA)(paired Wilcoxon tests); normalization for each cells to values before application. **D**, Phase plot analysis of AP generation before and after 50nM recombinant FGF13A extracellular application (paired Wilcoxon tests). **E-F**, Intrinsic excitability of mouse pyramidal neurons before and after 50nM synthetic ExonS extracellular application (2-way ANOVA test for AP firing rate; paired Wilcoxon test for rheobase). **G**, AP properties of mouse pyramidal neurons before and after 50nM synthetic ExonS extracellular application (paired Wilcoxon test). **H**, Phase plot analysis of AP generation before and after 50nM synthetic ExonS extracellular application (paired Wilcoxon tests). ns, non-significative; *, p<0.05; **, p<0.01; ***, p<0.001; ****, p<0.0001.

Collectively these data indicate that FGF13A alone can affect AP generation also at the soma, beyond the AIS, while the effects of LRRC37B overexpression (Figure 2E) are restricted to the AIS level. Finally, to relate these results to our molecular findings, we tested the impact of FGF13A ExonS, which is sufficient to bind to LRRC37B. We found that 50nM FGF13A ExonS elicited the same effects as full-length FGF13A (Figure 5E-H, S4F). In contrast, the same experiments performed with intracellular application of FGF13A at the same concentration (50nM) did not lead to detectable effects on neuronal excitability, except a small decrease in AP amplitude and sodium current (Figure S5), further indicating that at this concentration, FGF13A exerts its effects essentially at the extracellular level.

Altogether, these results indicate that FGF13A acts as a negative regulator of neuronal excitability through similar effects as LRRC37B, and suggest that LRRC37B acts as a regulator of FGF13A action by concentrating its effects at the level of the AIS.

### LRRC37B binds to the VGSC NAV beta subunit 1 (SCN1B)

As a complementary approach to identify binding partners of LRRC37B, we performed affinity purification of rat brain extracts using as a bait the recombinant ectodomain of LRRC37B fused to the Fc portion of human IgG (LRRC37B_ECTO-Fc) (Savas et al., 2014), followed by liquid chromatography tandem mass spectrometric (LC-MS/MS) analysis, compared to Fc alone. Several membrane proteins of the AIS were detected at low levels, including the NAV channel alpha subunit SCN1A, confirming our data, but also the regulatory NAV beta subunit SCN1B (Figure S6A). AIS proteins are resistant to detergents, which makes their extraction and identification challenging (Hamdan et al., 2020), hence the general low number of spectral counts.

Beta-subunits can directly bind to and positively regulate the trafficking and function of NAV channels, and are thereby critically involved in neuronal excitability and epilepsy (Namadurai et al., 2015). We therefore tested further their potential interactions with LRRC37B. Using heterologous co-expression in HEK cells, we found that the extracellular part of LRRC37B could be co-immunoprecipitated with SCN1B (Figure 6A-C). We next tested for the interaction of SNC1B with several LRRC37B deletion mutants. We found that the LB domain (amino acids 133-186 of LRRC37B), present only in LRRC37B and not in other LRRC37 paralogs, was necessary to mediate the binding between LRRC37B and SCN1B (Figure 6A-B). Consistent with the importance of the LB domain for SCN1B interaction, we detected no interaction between SCN1B and LRRC37A receptors that lack this domain, while macaque and chimpanzee LRRC37B receptors both displayed binding to SCN1B (Figure S6B). Overall, these data show that the LB domain is necessary and sufficient to mediate the binding between LRRC37B and SCN1B.

**Figure 6.**
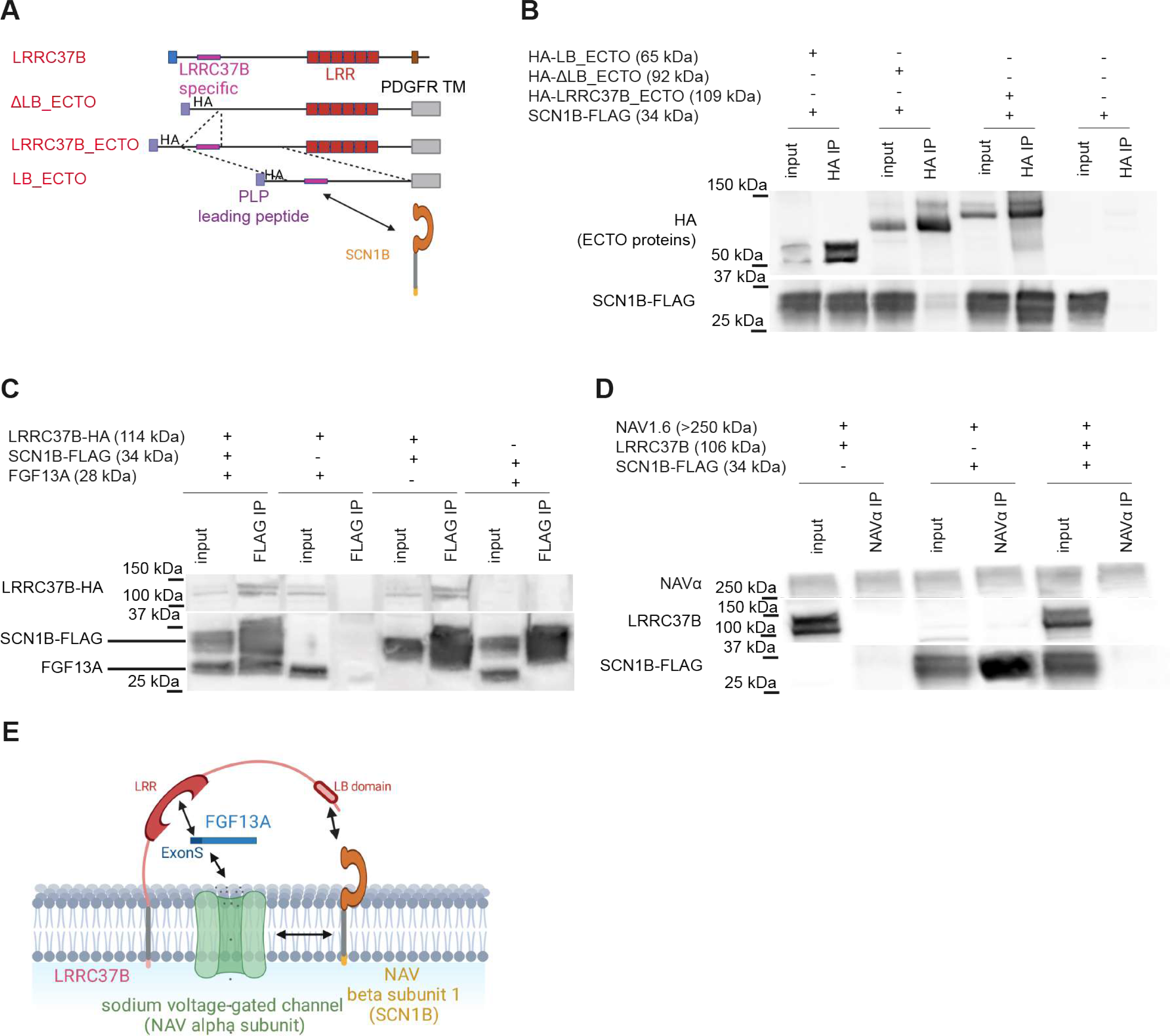
LRRC37B binds to SCN1B and prevents NAV1.6/SCN1B interaction. **A-B,** Immunoprecipitation of different protein lacking or carrying the LB specific domain of the extracellular domain of LRRC37B from HEK-293T cells co-transfected for the SCN1B cDNA. **C**, SCN1B immunoprecipitations from HEK-293T cells transfected for SCN1B +/- LRRC37B +/- FGF13A cDNAs. **D**, NAV1.6 immunoprecipitations from HEK- 293T cells transfected for SCN8A/NAV1.6 +/- SCN1B +/- LRRC37B cDNAs: SCN1B binds to NAV.6 which is altered in presence of LRRC37B. **E**, Schematic summary of the LRRC37B – SCN1B – FGF13A – NAV1.6 interaction.

Finally, we tested the interactions between LRRC37B and its binding partners FGF13A and SCN1B, as well as NAV1.6. We found that LRRC37B, SCN1B and FGF13A could be immunoprecipitated in the same complex, while SCN1B alone could not directly bind to FGF13A (Figure 6C). SCN1B was found to bind to NAV1.6, as expected (Namadurai et al., 2015), but this interaction was abolished when co-transfecting LRRC37B (Figure 6D). Given the known positive impact of SCN1B on NAV function (Namadurai et al., 2015), the ability of LRRC37B to compete for binding to NAV1.6 could lead to inhibitory influence on NAV function. On the other hand, as SCN1B promotes the targeting of NAVs to the AIS (O’Malley and Isom, 2015), the binding of SCN1B to LRRC37B could enable similarly the targeting of LRRC37B to the AIS, consistent with our observations that LRRC37A2, which lacks the B-domain and therefore does not bind to SCN1B (Figure S6B), is not targeted to the AIS (Figure S2D).

### LRRC37B regulates human cortical pyramidal neuron excitability

Could LRRC37B interactions with NAV1.6 regulators FGF13A and SCN1B be relevant in the human cerebral cortex? By examining scRNAseq databases (Bakken et al., 2021) we found that FGF13, SCN1B and SCN8A (NAV1.6) and LRRC37B, are co-expressed in most types of human cortical neurons, with FGF13A expressed at higher levels in interneurons, and LRRC37B/SCN1B in pyramidal neurons (Figure S7A). To test whether they can actually form the same complex as found *in vitro*, we performed co-immuno- precipitation experiments from human cortical tissue. Importantly, this revealed that NAV alpha subunits, LRRC37B, SCN1B, and FGF13A were found in the same complex (Figure 7A), thus confirming our *in vitro* findings in the human cortex *in vivo*. Finally, we tested the physiological relevance LRRC37B in human cortex by performing acute patch-clamp recordings on human *ex vivo* temporal cortex biopsies (Figure 7B, Table 1). Firstly, we confirmed that human upper layer CPNs display a lower excitability compared with mouse counterparts, as previously reported (Figure S7B) (Beaulieu-Laroche et al., 2018).

**Figure 7.**
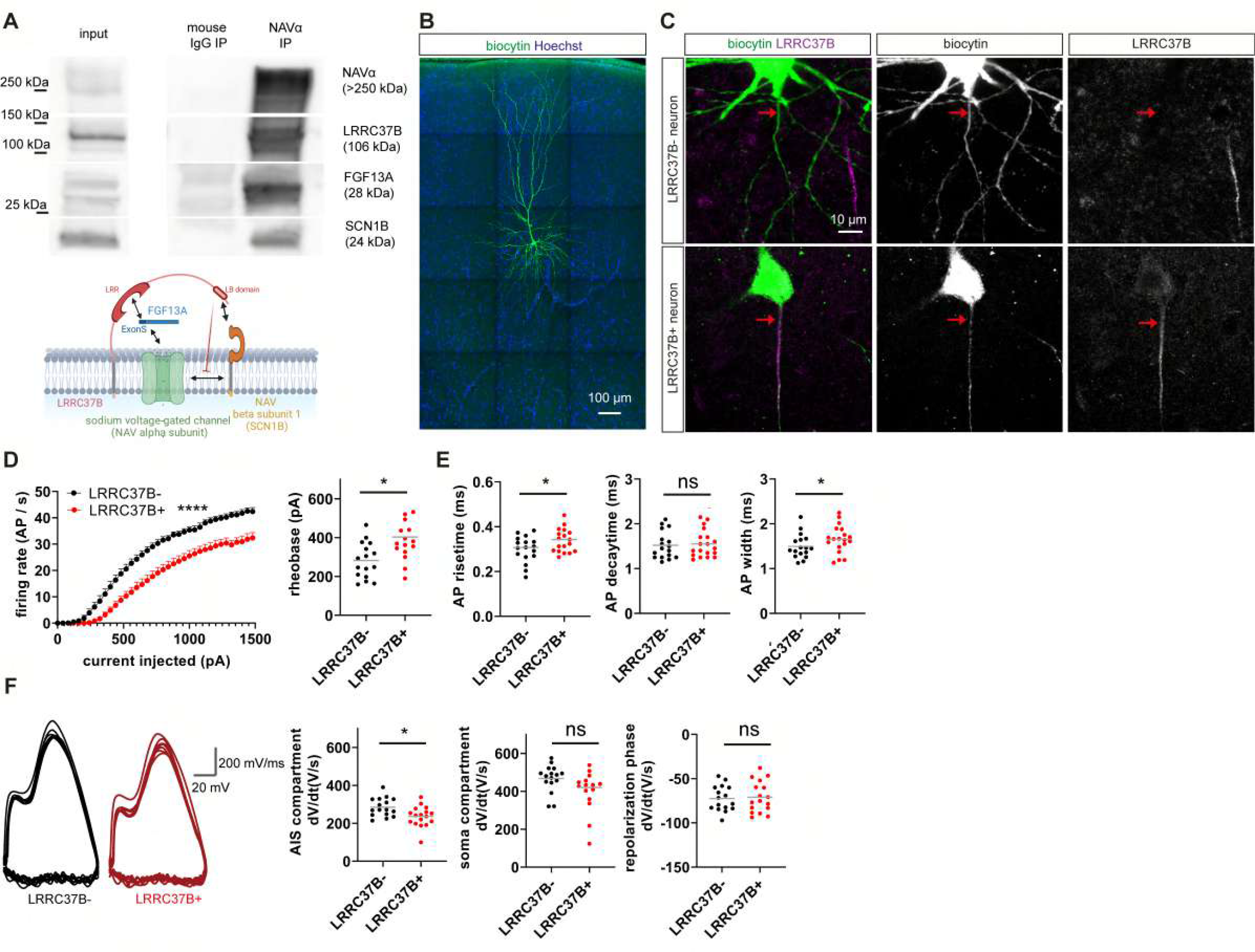
LRRC37B-positive human neurons display decreased neuronal excitability**. A**, NAVα immunoprecipitation from human cerebral cortex (25yo). **B-C**, Layer 2/3 human neurons filled by biocytin which enables to correlate their properties to the LRRC37B post-hoc detection (arrows)(B, 4yo; C, 61yo). **D-F**, Electrophysiological properties of LRRC37B-positive versus LRRC37B-negative neurons (2-way ANOVA test for the AP firing rate, Mann-Whitney test for other parameters)(4yo to 61yo cases). ns, non-significant; *, p<0.05; ****, p<0.0001.

Next, we tested whether the expression of LRRC37B in a subset of cortical neurons is relevant to their excitability. Since only a subset (∼40%) of human L2/3 PNs express LRRC37B (Figure 1D,E,I), we reasoned that the differential expression of LRRC37B would impact neuronal properties, specifically in this subset of cells. Therefore, during electrophysiological recordings, cells were filled with biocytin, followed by post-hoc LRRC37B immunostainings, to compare the properties of LRRC37B-positive vs LRRC37B-negative neurons (Figure 7C).

Remarkably, this revealed that LRRC37B-positive neurons display less excitability than LRRC37B-negative neurons, with a decreased AP firing rate, an increased rheobase and increased AP risetime and width (Figure 7D-E), while we observed no differences in input resistance, capacitance and sodium currents (Figure S7C-E). In addition, phase plot analysis of single AP revealed that LRRC37B-positive neurons display a decrease in the AIS compartment compared to LRRC37B-negative neurons (-17.2%), while other compartments remained unchanged (Figure 7F), thus mimicking the impact of LRC37B overexpressed in mouse CPNs (Figure 2E). Finally, as the length of the AIS has been implicated in the regulation of neuronal excitability and plasticity (Grubb and Burrone, 2010), we examined these parameters, but detected no differences in the AIS length of LRRC37B-positive AIS compared with LRRC37B- negative neurons (Figure S1F-H).

Overall these data indicate that expression of LRRC37B in pyramidal neurons can alter intrinsic excitability in the human cerebral cortex, in a way strikingly similar to what was observed in mouse neurons expressing LRRC37B, or treated with FGF13A.

## Discussion

Human brain evolution has been associated global changes in structure and function of the cerebral cortex, many of which have been linked to divergence in developmental mechanisms of cortical organogenesis (Libé-Philippot and Vanderhaeghen, 2021; Sousa et al., 2017). Distinctive evolutionary novelties of the human cerebral cortex also likely emerged from divergence at the level of individual neurons, leading to specialization in morphology and physiology.

Single cell profiling of human cortical neurons recently revealed that most cortical neuron types are shared between human and non-human primates, but that orthologous types display significant gene expression divergence (Bakken et al., 2021; Berg et al., 2021; Hodge et al., 2019). These likely account for some of the functional specializations of human CPNs, but the links between genomic and neuronal evolution have remained largely unknown.

Here we found that LRRC37B, encoded by a hominid-specific gene, acts as a species-specific modifier of human neuronal excitability, through direct interactions with FGF13A and SCN1B, two key regulators of voltage-gated ion channels. Interestingly, evolutionary changes targeting various classes of neuronal ion channels were previously reported in several species, including fish, flies and cephalopods, leading to species-specific neuronal features (Roberts et al., 2022).The expression levels of h-channels that modulate hyperpolarization-activated non-specific cation current, were also recently found to be expressed at higher levels in human than mouse cortical neurons (Kalmbach et al., 2018). Overall these data suggest that the modulation of membrane conductance properties is an important drive of neuronal evolution, for which LRRC37B gene duplication constitutes a striking example in our species.

Recent findings indicate that human neurons display specific distinctive functional properties not found in other species. In addition to increased synaptic connectivity (Charrier et al., 2012; Schmidt et al., 2021), function (Molnár et al., 2008, 2016), and plasticity (Szegedi et al., 2016; Testa-Silva et al., 2014), some classes of human CPNs display distinct biophysical and intrinsic properties, most of which converge towards decreased excitability (Beaulieu-Laroche et al., 2018, 2021; Deitcher et al., 2017; Eyal et al., 2016; Geirsdottir et al., 2019; Gidon et al., 2020; Kalmbach et al., 2018, 2021; Schwarz et al., 2019). Although some of these distinctive intrinsic properties may be linked in part to differences in the identity of the neurons analysed across species (Kalmbach et al., 2021; Moradi Chameh et al., 2021), they likely bear important evolutionary significance. The mechanisms underlying these differences have been mostly linked to specific properties of the dendrites, membrane capacitance, or ionic conductance. The involvement of the AIS (output modulation) had not been directly considered so far, even though some human cortical neurons were found to display distinct AP properties at the AIS (Testa-Silva et al., 2014).

Here we find that human CPNs display specialization of the AIS leading to decreased excitability, which we link to the selective expression of LRRC37B at this location.

Our data point to a model whereby LRRC37B binds to FGF13A and promotes its targeting to the AIS, which leads to the inhibitory modulation of the function of NAV channels, specifically at this subcellular site. As we find that FGF13A, acting as an extracellular ligand for NAV1.6, can decrease neuronal excitability more globally, also at somatic levels, it is tempting to speculate that it may act on other NAVα subunits located in other neuronal compartments, which remains to be explored.

Moreover we find that LRRC37B directly interacts with SCN1B, an important modulator of VGSC (Leterrier, 2016). Our data using heterologous expression indicate that LRRC37B could block the interaction of NAV1.6 with SCN1B, suggesting an additional mechanism by which LRRC37B could inhibit NAV channels. Interestingly, SCN1B was previously shown to interact with other transmembrane or extracellular proteins at the AIS (McEwen and Isom, 2004; Xiao et al., 1999) with potential impact on NAV channel function (Xiao et al., 1999). On the other hand, SCN1B promotes the targeting of NAV alpha subunits to the AIS (O’Malley and Isom, 2015), suggesting that it could similarly contribute to the specific AIS targeting of LRRC37B. Consistent with this hypothesis, LRRC37A paralog-encoded proteins are not targeted to the AIS.

In any case it remains to be determined whether the interaction of LRRC37B with SCN1B only leads to its selective targeting to the AIS, to additional inhibition of VGSC, or both. Another intriguing question that remains to address concerns the mechanisms that underlie the species-specific localization of LRRC37B at the AIS of human neurons, and whether this constitutes a uniquely human specialization. Indeed while LRRC37B proteins could not be detected in chimpanzee neurons in the samples that we probed, it remains to be determined whether it is expressed at other stages, ages or brain areas.

We identify FGF13A as an extracellular ligand for LRRC37B and NAV1.6. This is unexpected since FHF non-canonical FGFs (FGF11-13) were reported to encode non-secreted proteins, based on the absence of a predicted signal peptide (Smallwood et al., 1996). Notably however, proteins lacking N-terminal signal peptides, among them FGF2, can still be secreted trough non-canonical mechanisms (Rabouille, 2017). Inhibitory effects of intracellular FGF13A on NAVα subunits were suggested previously (Abrams et al., 2020; Barbosa et al., 2017; Dover et al., 2010; Effraim et al., 2019, 2022; Yang et al., 2016). However, these studies either used FGF13A transfection or very high (1mM) intracellular concentrations of FGF13 (Barbosa et al., 2017; Dover et al., 2010), while at the more physiological (nM) concentrations used in our study, no effect is detectable following intracellular addition of FGF13A, except a mild reduction of sodium current intensities and AP amplitude. Our data thus indicate that FGF13A acts mostly extracellularly to decrease neuronal excitability, while not excluding that other isoforms of FGF13 or its core domain could act intracellularly (Barbosa et al., 2017; Effraim et al., 2019).

FGF13-like genes are found in all metazoans and are the ancestors of all vertebrate FGF proteins (Itoh and Ornitz, 2008). In addition, the ExonS of FGF13A, which mediates secretion of the ligand and its binding to LRRC37 proteins, is conserved between species and found in the other paralogs of the FHF family (Fry et al., 2021), suggesting that other members of the family constitute secreted ligands.

Given the extracellular location of FGF13A, an important open question is the cellular origin of the ligand. FGF13 has been found at the level of the AIS of pyramidal neurons in the hippocampus (Pablo et al., 2016) and we find it similarly located in cortical CPNs, suggesting that the regulation by FGF13A could be autocrine, at least in part. Interestingly, *FGF13* is also expressed in chandelier interneurons (ChC), which are key regulators of CPN excitability through their axo-axonic inhibitory synapses targeting the AIS of CPNs (Favuzzi et al., 2019). FGF13A could thus be secreted by ChC and thereby modulate NAV channel activity, and this interaction could be enhanced by the presence of LRRC37B at the AIS. Moreover and interestingly, *FGF13* was previously found to be necessary for the level of innervation of the AIS of mouse CPNs (Favuzzi et al., 2019). While we did not detect any effect of LRRC37B on the patterns of inhibitory innervation of the AIS, it will be interesting to determine whether LRRC37B and FGF13A modulate other aspects of CPN-ChC interactions. From scRNAseq data, LRRC37B is also expressed in all classes of interneurons, although at lower levels than in CPNs, and could also act there to modulate neuronal excitability, in an autocrine or paracrine fashion. Finally, an intriguing finding is that LRRC37B is expressed in only in a subset of all cortical neuron subtypes, independently of their identity. This suggests that the levels of expression of this receptor could depend on cell context and state, such as the levels of neuronal activity, which will be important to explore.

The discovery of LRRC37B as a co-receptor for FGF13A and SCN1B regulating NAV channels at the AIS has potentially important implications for brain diseases, in particular epilepsy. Patients displaying specific mutations at the level of the ExonS of FGF13A, the site of interaction to LRRC37B, are affected by epilepsy (Fry et al., 2021; Narayanan et al., 2022). Similarly, SCN1B, the other interactor of LRRC37B that we identified, is mutated in severe forms of epilepsy, resulting mostly in deregulation of NAV channels (Devinsky et al., 2018; Wimmer et al., 2010). It will be important to test whether and how pathogenic mutations in *FGF13* and *SCN1B* genes impact on their ability to interact with LRRC37B, as well as search for potential pathogenic mutations or pathogenic copy-number variations of *LRRC37* genes. In this respect our finding that the copy number of LRRC37B is nearly fixed in the normal human population strongly suggests its importance for normal human physiology. While no evidence for germ-line mutations can be found in databases, the possibility of somatic mutations (Zhang and Vijg, 2018) is an interesting possibility that remains to be explored.

On the other hand, given its strong impact on neuronal intrinsic properties, LRRC37B is likely to play important roles during normal cortical circuit function. The potential consequences of human-specific changes in neuronal properties on circuit function remain poorly known, but the identification of LRRC37B as a human-specific modulator of neuronal excitability opens new avenues to study this experimentally. LRRC37B could affect information processing through neuronal gain modulation, by which neurons adapt to changing inputs (Ferguson and Cardin, 2020) or amount of information transfer (Testa-Silva et al., 2014). Indeed, while neuronal gain modulation relies largely on the regulation of synaptic inputs, it can also be modified by output modulation, including at the level of the AIS (Debanne et al., 2019). Moreover, neuronal plasticity and learning involve modulation of the neuronal gain including the output site (Ferguson and Cardin, 2020; Jamann et al., 2021). The modulation of excitability by LRRC37B could thus influence neuronal information processing and plasticity, at least in part through gain modulation and amount of information transfer, and thereby contribute to human-specific properties of cortical circuits.

While LRRC37B is likely to be unique among LRRC37 receptors for its ability to bind to SCN1B, as it is mediated by the LB domain found only in this LRRC37 paralog, other LRRC37A-type receptors are also expressed in cortical neurons and have the ability to bind FGF13A. It will be interesting to test their function, which could involve modulation of different types of voltage-gated ion channels at other subcellular compartments, such as cell body and dendrites. Similarly, it will be important to test whether and how *LRRC37B*, as well as other members of the *LRRC37* family, are expressed in other brain areas and tissues, where they may act as human-specific modifiers of neuronal excitability or other cellular features selected throughout recent evolution. For instance, LRRC37A2 was recently found to interact with alpha-synuclein in astrocytes (Bowles et al., 2022).

Several HS gene duplicates were linked previously to the control of corticogenesis (Charrier et al., 2012; Fiddes et al., 2018; Florio et al., 2015; Van Heurck et al., 2022; Schmidt et al., 2021; Suzuki et al., 2018), but so far none have been linked to specific functional features of human neurons. While *LRRC37B* is the first example of a gene duplicated controlling adult human neuron function, another set of human-specific genes, *SRGAP2B/C*, were found to control the development of cortical synapses (Charrier et al., 2012). A mouse model displaying expression of *SRGAP2C* in pyramidal neurons leads to hyper-connectivity of the mouse neurons and as a result enhanced processing ability and improved behavioural performance of the transgenic mice (Schmidt et al., 2021). It will be interesting to test whether and how the emergence of *LRRC37* and *SRGAP2* genes evolved together or separately to control excitability and connectivity of human cortical neurons.

In conclusion we identified a new molecular pathway regulating membrane excitability at the level of the AIS in human cortical neurons, targeting the highly conserved NAVs, but modulated in a species-specific and multimodal fashion by LRRC37B and its interactors. These data provide unexpected molecular links between the subcellular regulation of neuronal excitability and human brain evolution, with important implications for our understanding of human brain function, and diseases.

## Acknowledgments

We thank members of the PV lab and CBD for helpful discussions and precious help. We thank patients for their participation in the study and Anaïs van Hoylandt for assistance. We thank Juan Burrone (King’s College, United Kingdom) and Franck Polleux (Columbia University, United States) for excellent advice and for sharing precious reagents. We thank Michele Giugliano (International School for Advanced Studies, Italy) and Linda van Aelst (Cold Spring Harbour Laboratory, USA) for excellent advice. We thank the VIB Proteomics Core (Gent, Belgium) led by Francis Impens for the LC-MS/MS analyses. We thank Bart de Strooper lab (CBD) for sharing its chemiluminescence imaging system. We thank Patrick Verstreken lab (CBD) for sharing its absorbance plate reader. Some of the images were acquired on Zeiss LSM 880 and Abberior STED systems supported by Hercules AKUL/15/37_GOH1816N and FWO G.0929.15 to Pieter Vanden Berghe, KU Leuven; we thank Pieter Vanden Berghe and Tobie Martens for trainings and technical advice. The authors gratefully acknowledge the VIB Bio Imaging Core for their support & assistance in this work. Diagrams were created with BioRender.com. This work was funded by Grants of the European Research Council (GENDEVOCORTEX), the EOS Programme, ERANET NEURON, the Belgian FWO and FRS/FNRS, the Generet Foundation, and the Belgian Queen Elizabeth Foundation (to PV) and in part by the RWJ Foundation (#74260), the National Science Foundation award (#1755189) (to DC) and the National Institutes of Health (DP2MH119424) (to DS and MYD). B. L.-P. was supported by a postdoctoral Fellowship of the FWO (12V1219N).

## Author contributions

Conceptualization and Methodology, BLP, PV; Investigation, PV.; Copy-numbers analysis, DS, MYD; Electrophysiological recording and analysis, KW, IV, AL; In utero electroporations, SB, BLP; Molecular cloning assistance, AL, VG; Cryosections, AB, BLP; Protein pull down experiment, KMV, JdW; Recruitment and informed consent of human subjects, surgical resection of human cortical tissue, TT; Human biopsy processing assistance, AL, HN; Elisa assay, TWB, DC; All other experimental work and analyses, BLP, AL; Formal Analysis,; Writing, PV; Funding acquisition, PV; Resources, PV; Supervision, JdW, PV. Competing interests: Authors declare no competing interests. Data and materials availability: All data are available in the manuscript or the supplementary materials. All materials are available upon request from PV.

## Supplementary Materials

### Supplementary Figure Legends

**Figure S1.**
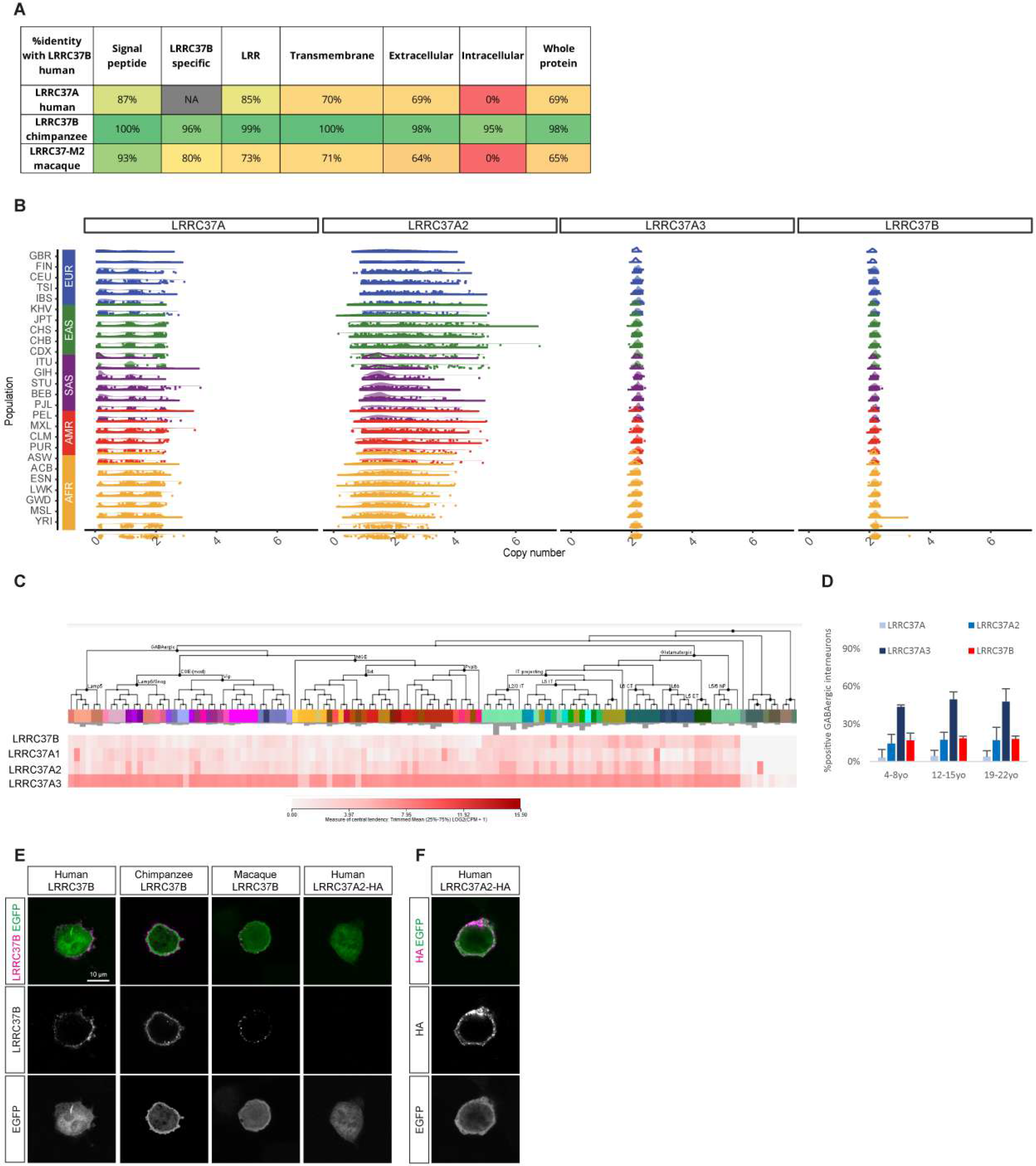
(related to Figure 1). Structure and expression of LRRC37B. **A,** Percentage of similarity of amino acid sequence between human *LRRC37B* and chimpanzee and macaque orthologs as well as *LRRC37A* paralog. **B,** Copy numbers of *LRRC37* encoding genes in modern human populations (EUR, European; EAST, East Asian; SAS, South Asian; AMR, American; AFR, African; subpopulations defined by the 1000 Genome Project). **C**, *LRRC37* encoding transcripts detection in human cortical neurons (from Allen Brain Atlas and (Bakken et al., 2021)). **D**, Proportion of GABAergic interneurons at different ages with *LRRC37* transcripts detection from human postnatal cortical samples (yo: years old, from (Velmeshev et al., 2019), mean + SEM). **E**, Live immunostaining for LRRC37B (see Methods) on HEK-293T cells transfected for human, chimpanzee and macaque LRRC37B as well as human LRRC37A2-HA (single focal plan). **F**, Immunostaining after fixation and permeabilization for HA on HEK-293T cells transfected for LRRC37A2-HA (single focal plan).

**Figure S2.**
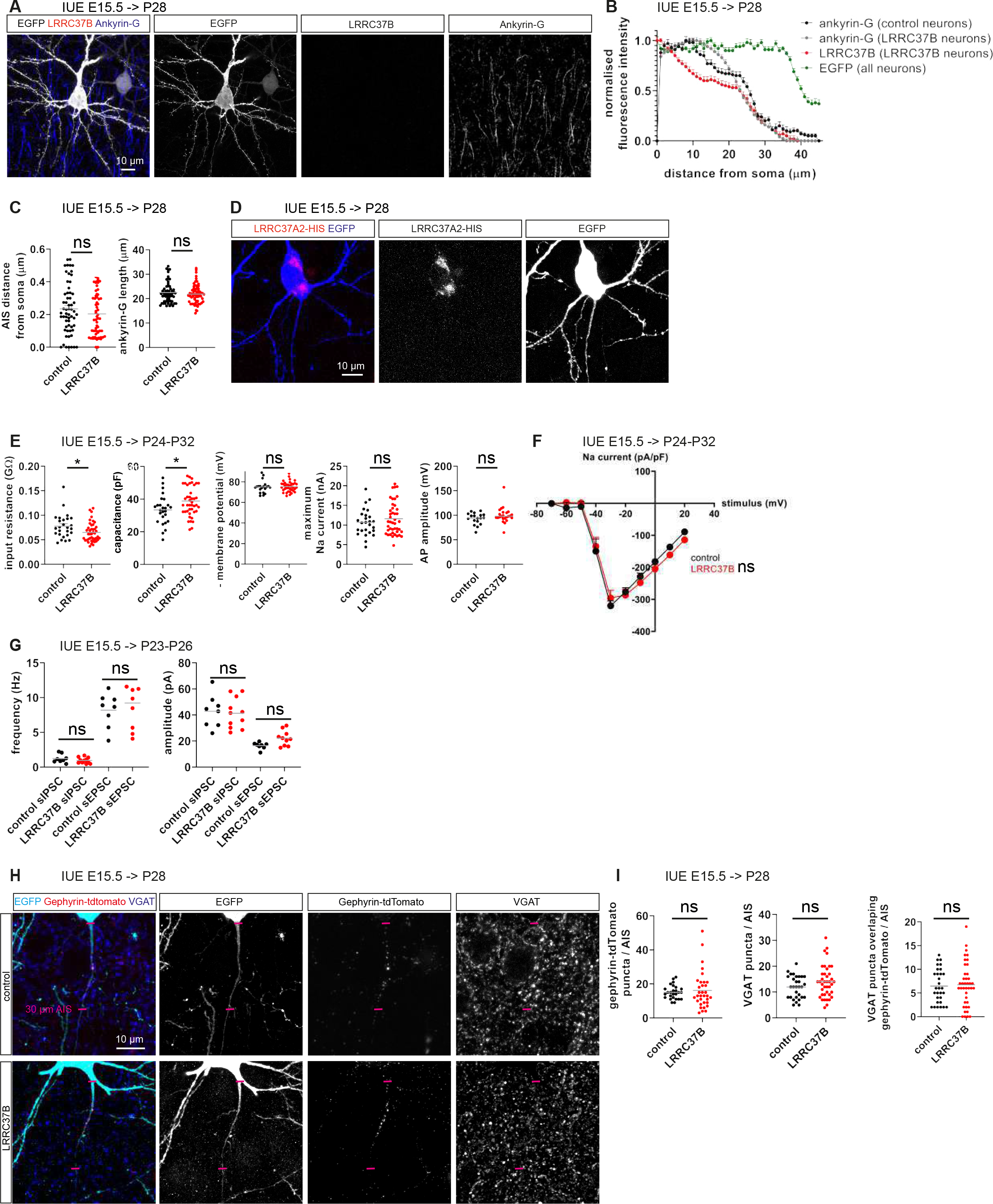
(related to Figure 2). LRRC37B effects on functional and synaptic properties. **A,** Immunodetection of LRRC37B and Ankyrin-G in the mouse cerebral cortex after transfection of EGFP cDNA. **B-C,** LRRC37B colocalizes with Ankyrin-G in mouse neurons transfected for LRRC37B and EGFP with no differences in the AIS length or position of LRRC37B/EGFP neurons compared to EGFP control neurons (Mann-Whitney tests). **D**, Immunodetection of LRRC37A2-HIS in the mouse cerebral cortex after transfection of LRRC37A2-HIS and EGFP cDNAs. **E-F**, Electrophysiological parameters related to Figure 2B-D (2-way ANOVA for sodium currents, Mann-Whitney tests for others). **G**, Excitatory and inhibitory postsynaptic potentials (E/I PSP) frequency and amplitude in LRRC37B/EGFP versus EGFP control neurons (Mann-Whitney tests). **H-I**, Quantification of VGAT puncta, Gephyrin-tdTomato puncta and VGAT/Gephyrin-tdTomato puncta in mouse neurons in utero electroporated for cDNAs coding for LRRC37B and EGFP or EGFP alone as well as Gephyrin-tdTomato (Mann-Whitney tests). ns, non-significative; *, p<0.05.

**Figure S3.**
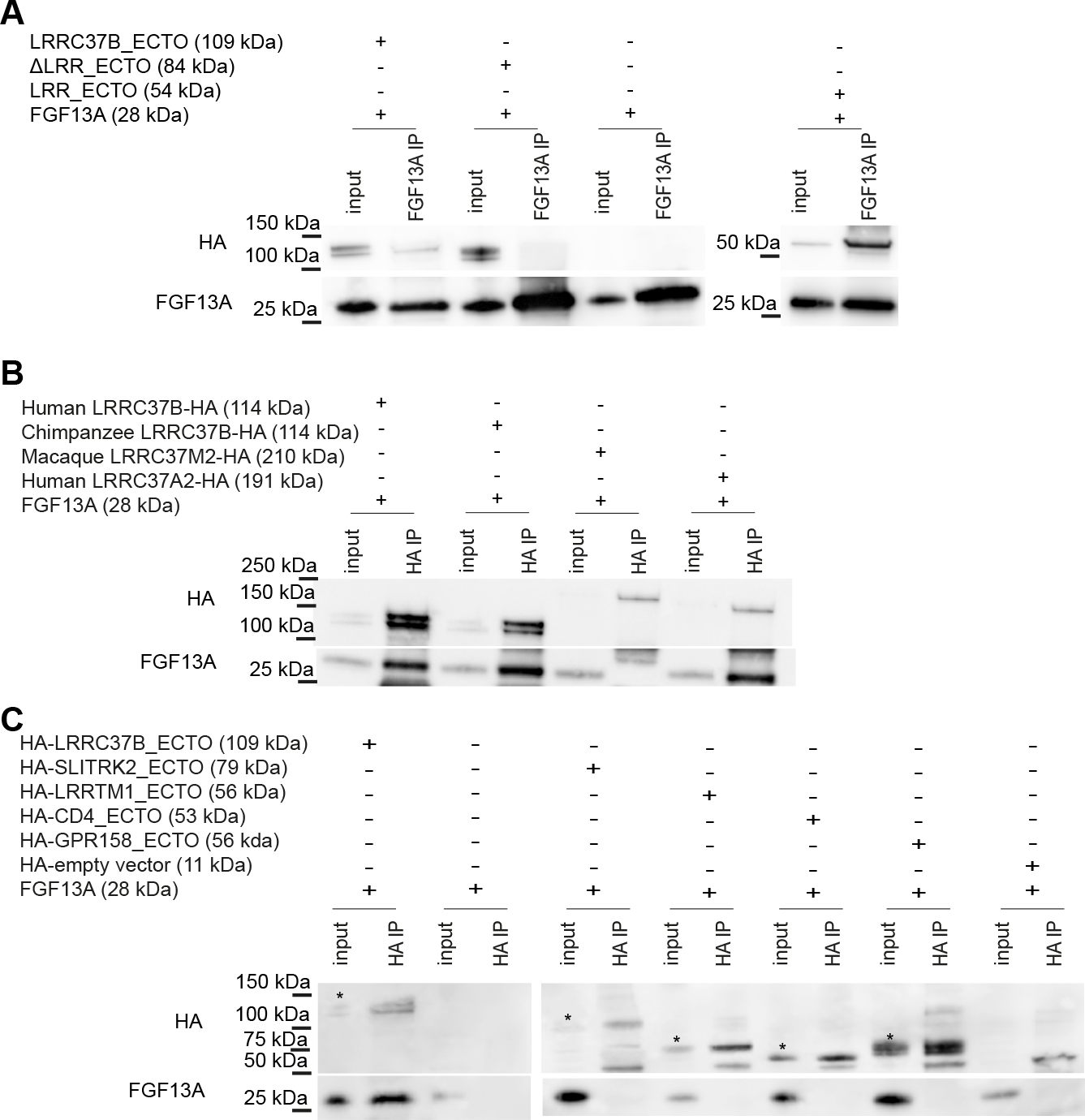
(related to Figure 3). LRRC37 proteins are co-receptors for FGF13A. **A,** FGF13A co-immunoprecipitates the LRRC37B extracellular part (LRRC37B_ECTO) as well as its leucin-rich repeats (LRR) but not the extracellular part devoid of the LRR from transfected HEK-293T cells. **B**, Immunoprecipitations of human, chimpanzee, macaque LRRC37B proteins or human LRRC37A2 protein from HEK-293T cells transfected for their cDNA and FGF13A cDNA: all LRRC37 proteins except the macaque LRRC37B binds to FGF13A. **C**, Immunoprecipitations of LRRC37B_ECTO and other transmembrane proteins (extracellular domain fused at the N-terminal with the prolactin leader peptide and an HA tag, and at the C-terminal with the transmembrane domain of PDGF-R) from HEK-293T cells transfected for their cDNA and FGF13A cDNA; stars indicate the transmembrane protein in the input.

**Figure S4.**
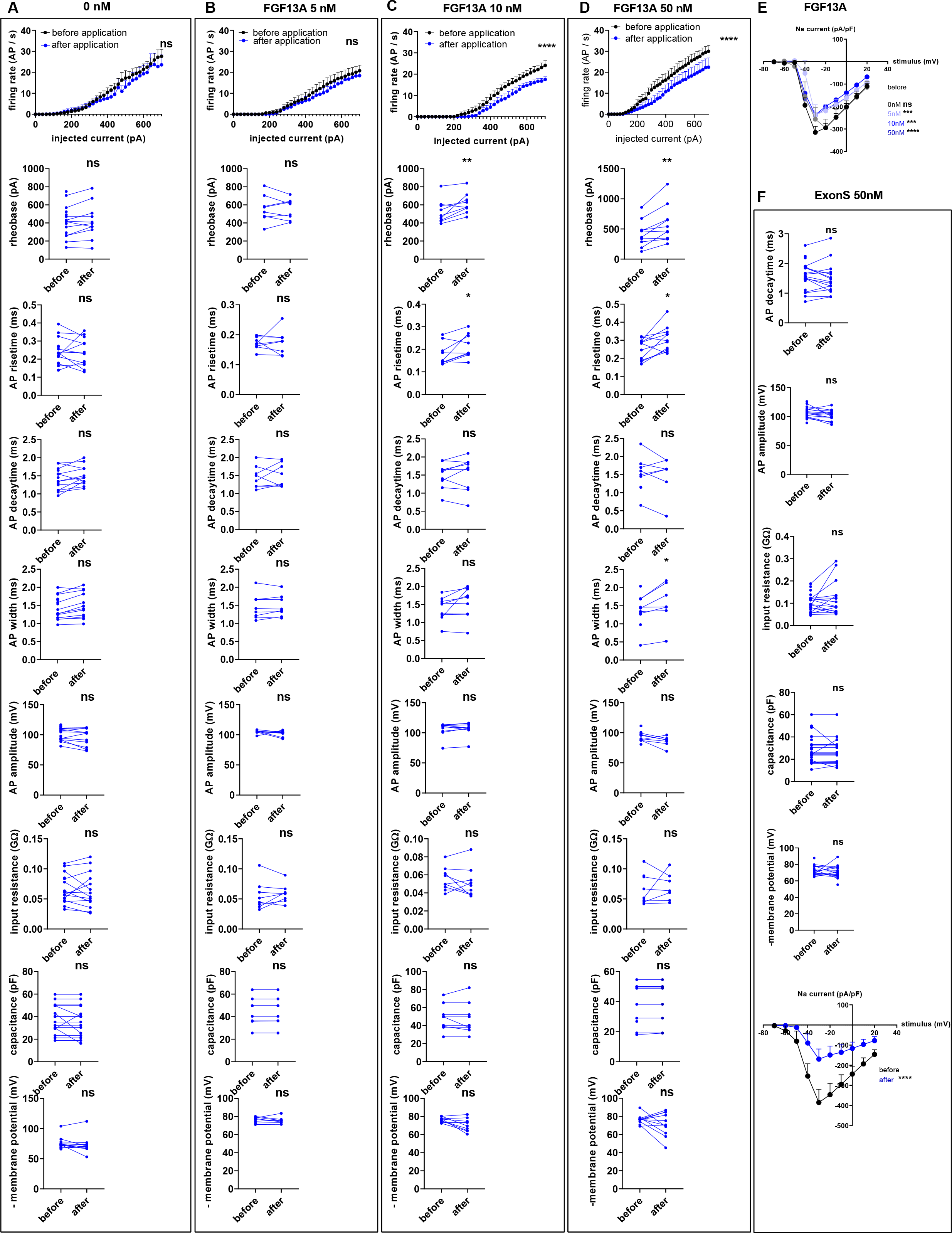
(related to Figure 5). FGF13A and its ExonS act extracellularly on neuronal excitability. **A-D,** Electrophysiological properties complementary to Figure 45A-C, with recombinant FGF13A extracellular application on mouse cortical sections at 0nM, 5nM, 10nM and 50nM (2-way ANOVA tests for AP firing rate, paired Wilcoxon test for others). **E**, Dose-response (0-50nM) of sodium currents of mouse cortical neurons with recombinant FGF13A extracellular application. **F**, Electrophysiological properties complementary to Figure 5E-G, with synthetic ExonS extracellular application on mouse cortical sections at 50nM (2-way ANOVA tests for AP firing rate & sodium currents, paired Wilcoxon tests for others). ns, non-significant; *, p<0.05; **, p<0.01; ***, p<0.001; ****, p<0.0001.

**Figure S5.**
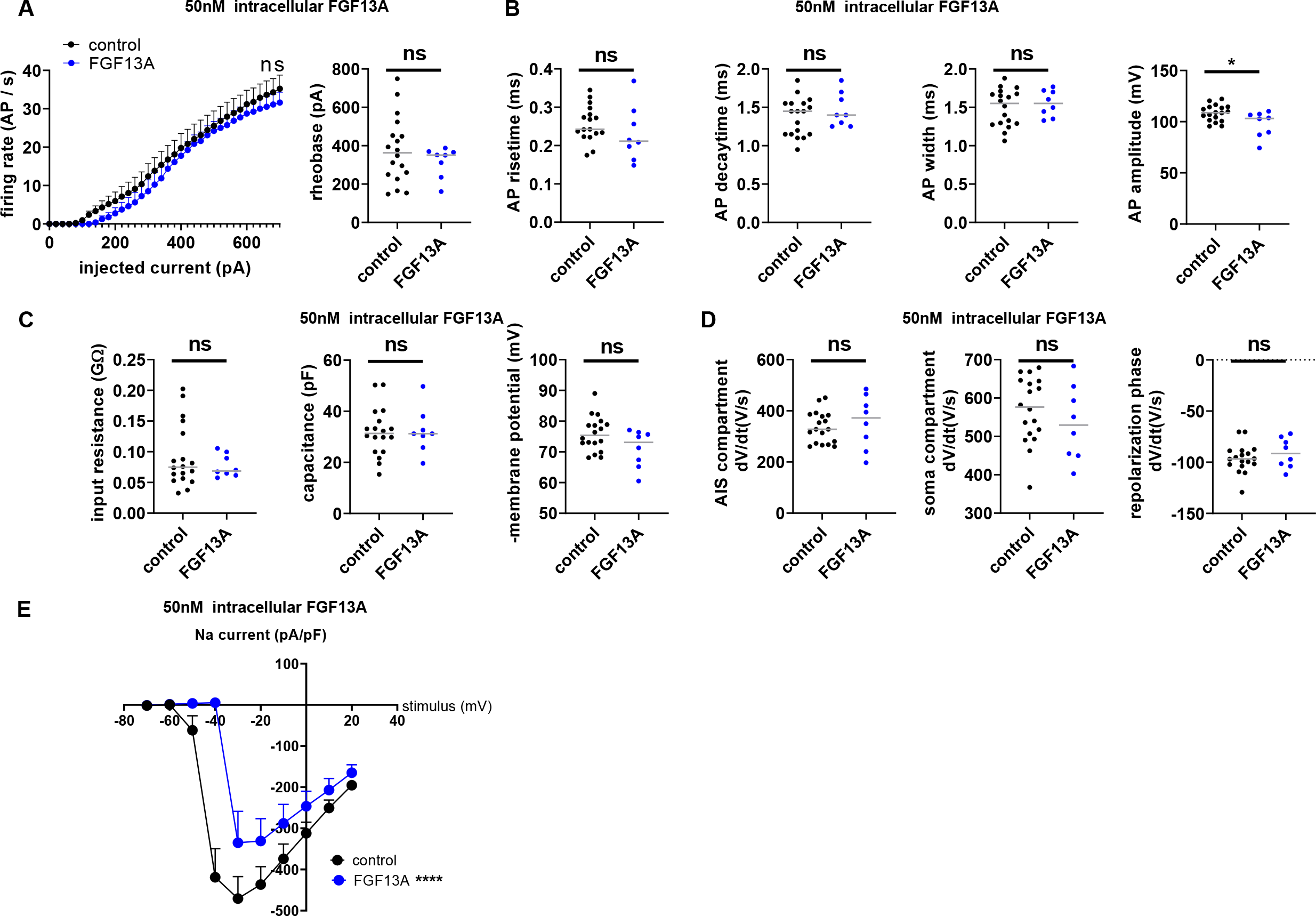
(related to Figure 5). FGF13A doesn’t act intracellularly on neuronal excitability. **A,** Cell intrinsic excitability of mouse neurons with 50nM intracellular application of recombinant FGF13A (2-way ANOVA test for AP firing rate, Mann-Whitney test for rheobase). **B**, AP properties of mouse neurons with 50nM intracellular application of recombinant FGF13A (Mann-Whitney tests). **C**, Electrophysiological properties of mouse neurons with 50nM intracellular application of recombinant FGF13A (Mann-Whitney tests)**. D,** Phase plot analysis of single APs of mouse neurons with 50nM intracellular application of recombinant FGF13A (Mann-Whitney tests)**. E,** Sodium currents of mouse neurons with 50nM intracellular application of recombinant FGF13A (2-way ANOVA). ns, non-significant; *, p<0.05; ****, p<0.0001.

**Figure S6.**
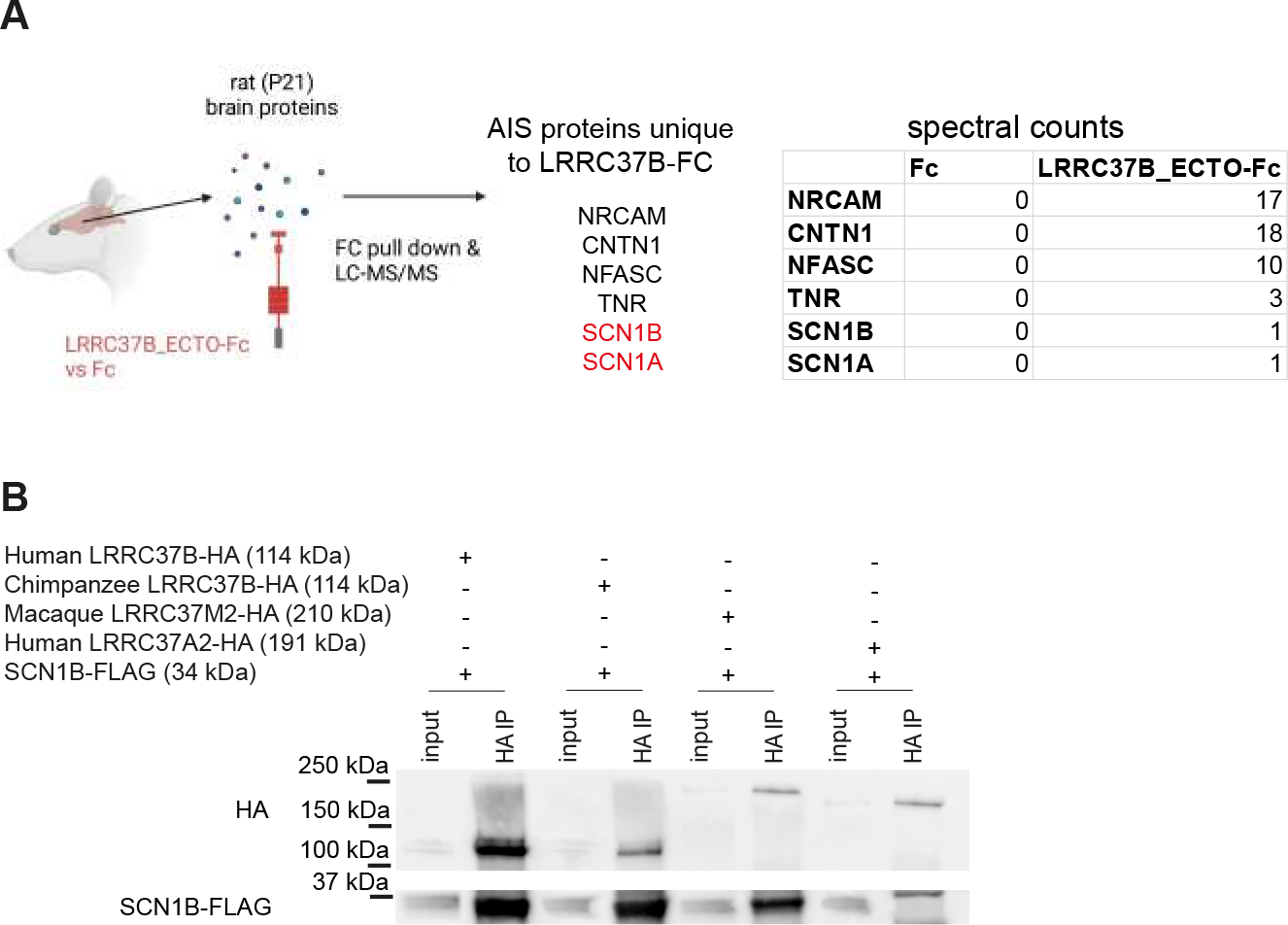
(related to Figure 6): LRRC37B binds to SCN1B through its specific domain. **A,** LRRC37B-Fc pull down from rat cortical extracts revealed its putative interaction with AIS proteins. **B,** Immunoprecipitations of human, chimpanzee, macaque LRRC37B proteins or human LRRC37A2 protein from HEK-293T cells transfected for their cDNA and SCN1B cDNA: all LRRC37B proteins binds to SCN1B but not LRRC37A2.

**Figure S7.**
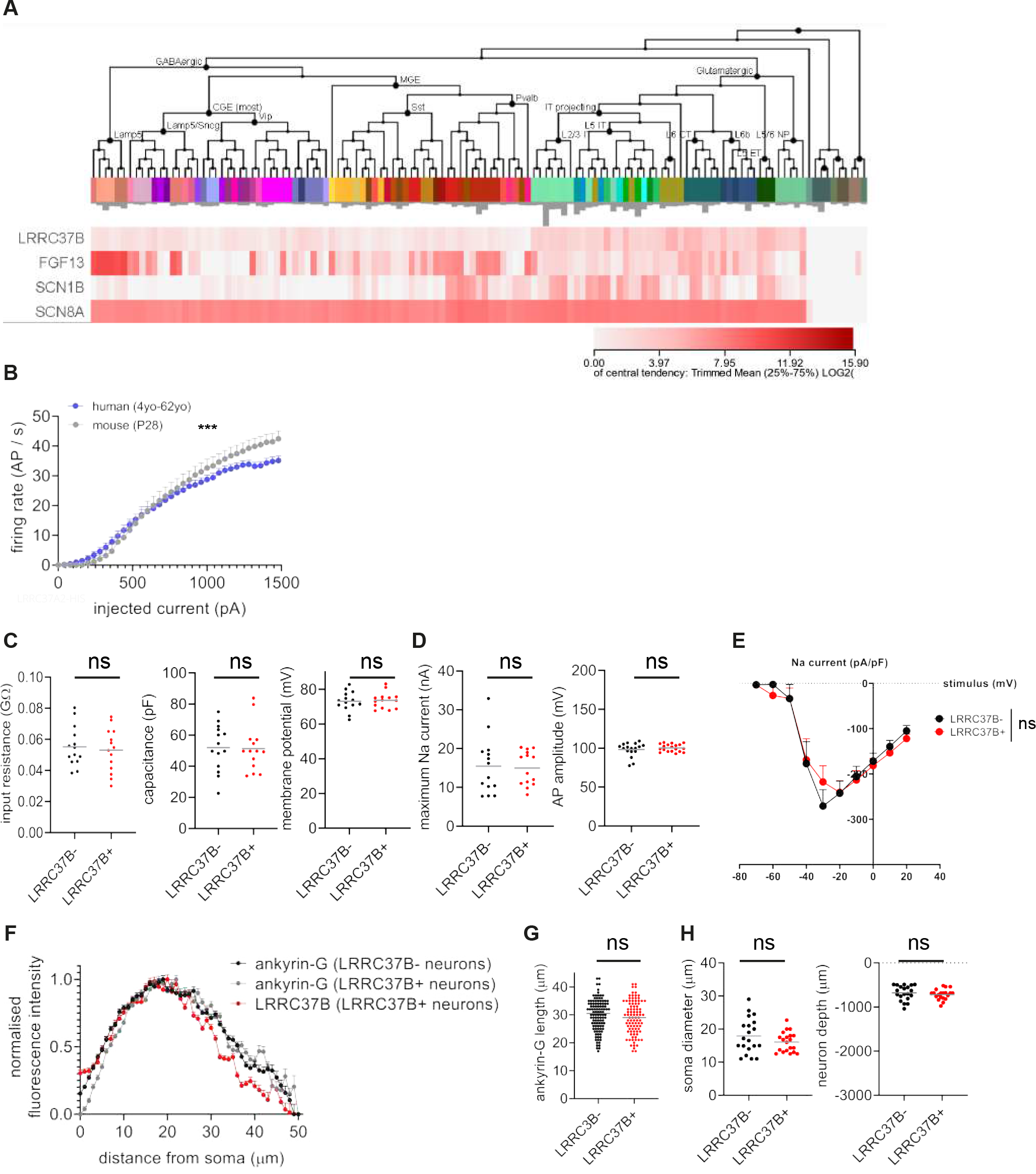
(related to Figure 7). LRRC37B function in human neurons**. A**, *LRRC37B*, *FGF13*, *SCN1B* and *SCN8A* transcripts detection in human cortical cells (from Allen Brain Atlas and (Bakken et al., 2021)). **B**, Mouse pyramidal neurons display a higher excitability than human neurons (data for human neurons also shown in Figure 67D; 2-way ANOVA test; ***, p<0.001). **C-E**, Electrophysiological properties of human neurons complementary to Figure 7D-F (2-way ANOVA for sodium currents, Mann-Whitney tests for others). **F**, Morphological properties of human neurons complementary to Figure 1G (Mann-Whitney test). **H**, Morphological properties of human neurons complementary to Figure 7B-C (Mann-Whitney test). ns, non-significant

### Supplementary Tables

**Table 1:** Human samples Autopsies have been used for immunostainings, biopsies have been used for immunostainings, electrophysiological recordings and immunoprecipitations

| Age | Gender | Origin | Autopsy/<br>Biopsy | Diagnosis | Cortical area |
| --- | --- | --- | --- | --- | --- |
| **3 months** | F | UK Brain Bank | autopsy | accidental asphyxia | temporal |
| **4 months** | M | UK Brain Bank | autopsy | coronary heart disease | temporal |
| **21 months** | F | UK Brain Bank | autopsy | coronary heart disease | temporal |
| **2 years** | M | UZ Leuven | biopsy | temporal lobe epilepsy | temporal |
| **4 years** | F | UZ Leuven | biopsy | temporal lobe epilepsy | temporal |
| **7 years** | M | UZ Leuven | biopsy | focal cortical displasia | temporal |
| **7 years** | M | UZ Leuven | biopsy | temporal lobe epilepsy | temporal |
| **11 years** | M | UZ Leuven | biopsy | epilepsy | fronto-insular |
| **14 years** | F | UZ Leuven | biopsy | focal cortical displasia | temporo-parietal |
| **14 years** | M | UZ Leuven | biopsy | epilepsy | frontal |
| **16 years** | M | UK Brain Bank | autopsy | sudden accident | temporal |
| **16 years** | M | UK Brain Bank | autopsy | suspension by ligature | occipital |
| **18 years** | M | UZ Leuven | biopsy | temporal lobe epilepsy | temporal |
| **22 years** | F | UZ Leuven | biopsy | temporal lobe epilepsy | temporal |
| **24 years** | F | Dutch Brain Bank | autopsy | donor control (no degeneration) | superior occipital + superior parietal |
| **25 years** | F | UZ Leuven | biopsy | temporal lobe epilepsy | temporal |
| **27 years** | F | UZ Leuven | biopsy | focal cortical displasia | temporal |
| **29 years** | M | UZ Leuven | biopsy | temporal lobe epilepsy | temporal |
| **38 years** | F | UZ Leuven | biopsy | temporal lobe epilepsy | temporal |
| **45 years** | M | Dutch Brain Bank | autopsy | donor control (no degeneration) | superior parietal |
| **45 years** | M | Dutch Brain Bank | autopsy | donor control (no degeneration) | lateral occipital |
| **46 years** | F | UZ Leuven | biopsy | temporal lobe epilepsy | temporal |
| **48 years** | M | UZ Leuven | biopsy | temporal lobe epilepsy | temporal |
| **62 years** | M | UZ Leuven | biopsy | temporal lobe epilepsy | temporal |

**Table 2:**
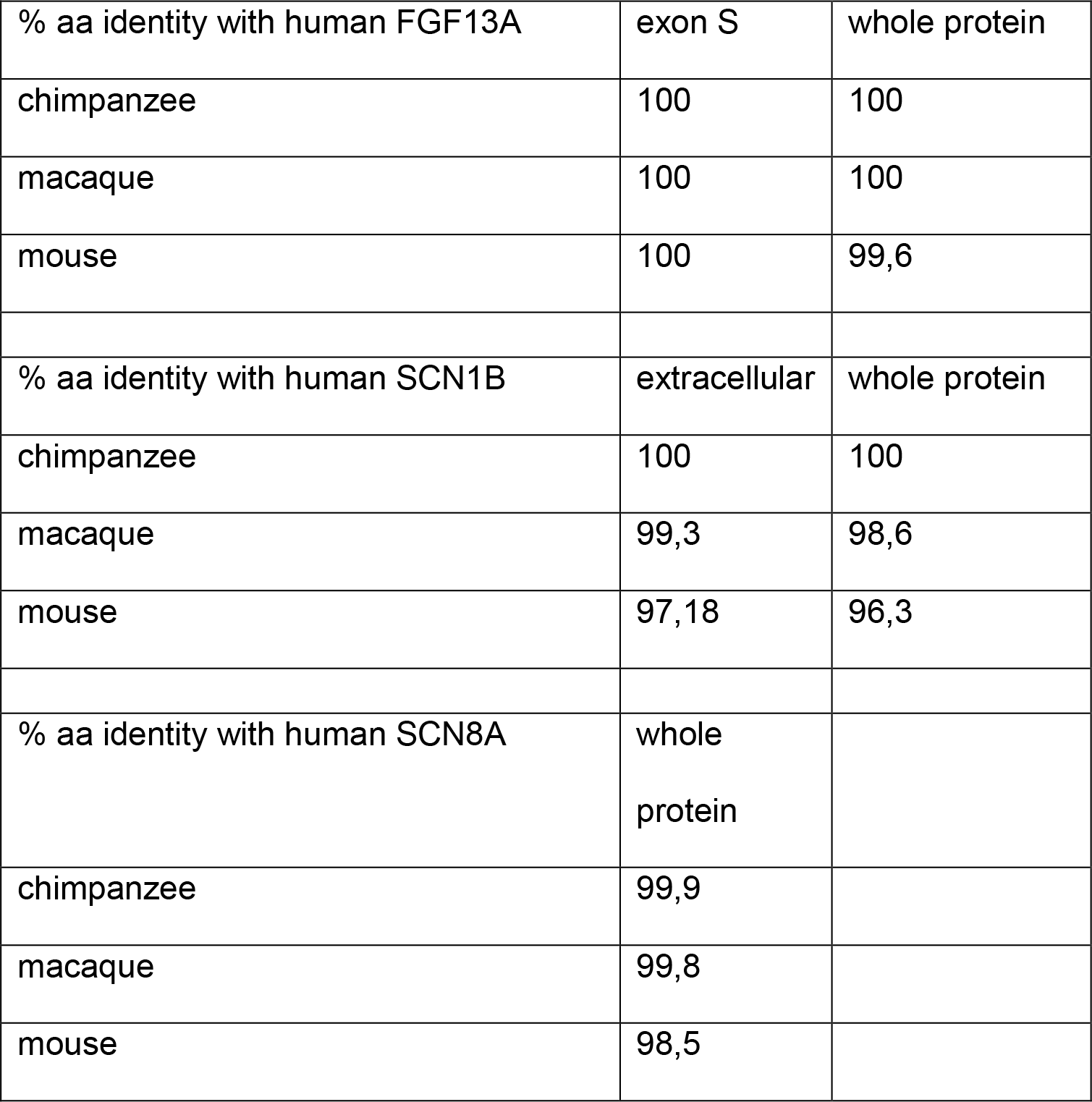
Similarities of FGF13A, SCN1B and SCN8A across species

**Table 3:**
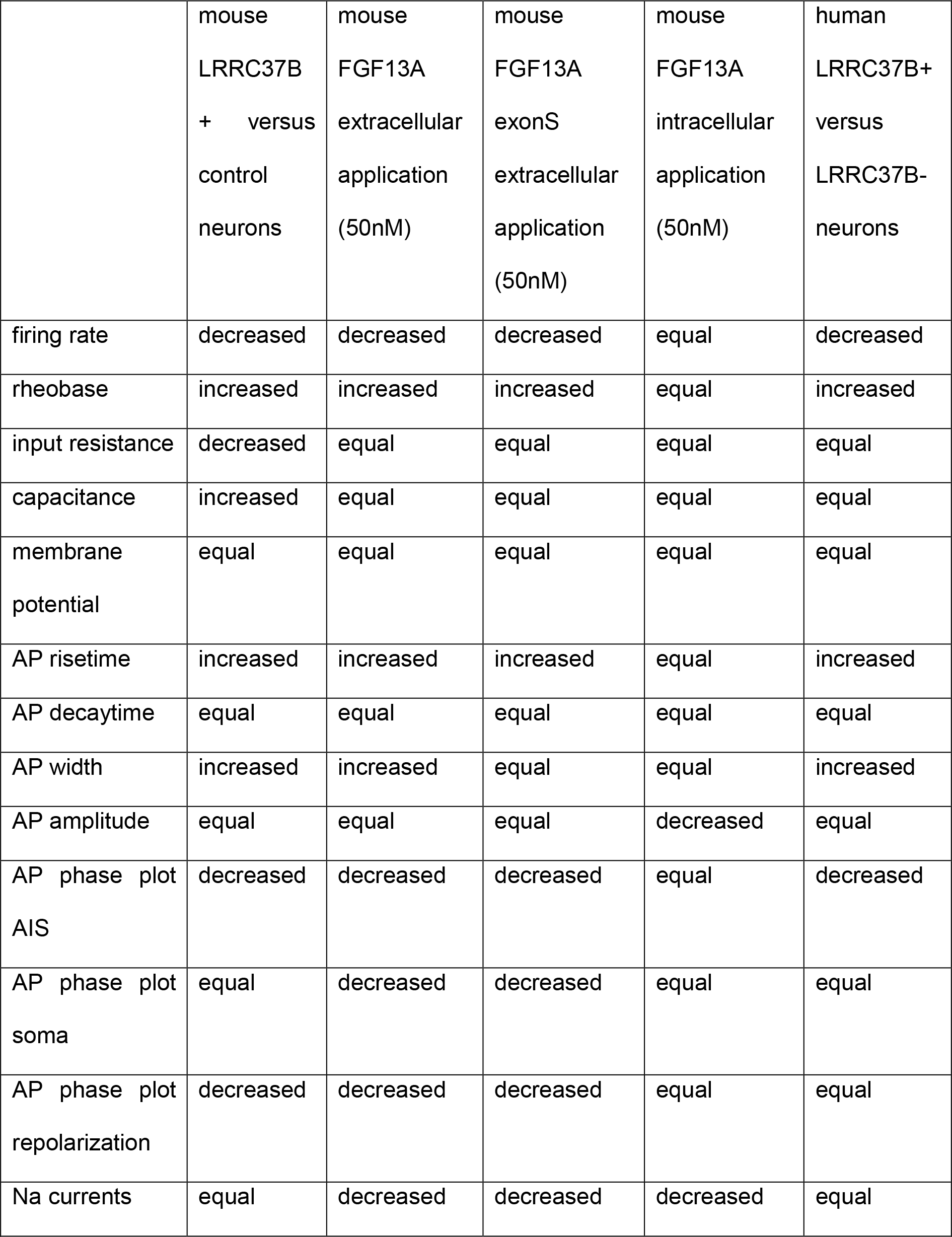
Summary of electrophysiological phenotypes between experiments

**Table 4:** Summary of species differences

|  |  | mouse | macaque | chimpanzee | human | specificity |
| --- | --- | --- | --- | --- | --- | --- |
| genome | B-type | no | yes | yes | yes | simian |
|  | LRRC37B | no | no | yes | yes | hominid |
| Transcripts<br>detection | LRRC37<br>genes | no | yes | ? | yes | simian |
|  | B-type | not<br>applicable | yes | ? | yes | simian |
|  | LRRC37B | not<br>applicable | not<br>applicable | ? | yes | hominid or<br>human |
| AIS<br>localization | LRRC37B | not<br>applicable | no | no | yes | human |
| protein<br>properties | LRRC37<br>proteins -<br>FGF13A | yes<br>(prediction<br>to the A<br>type) | yes<br>(prediction<br>to the A<br>type) | yes | yes | at least<br>mammal |
|  | B-type -<br>FGF13A | no | no | yes | yes | hominid |
|  | LRRC37<br>proteins -<br>SCN1B | no<br>(prediction<br>to the A<br>type) | yes | yes | yes | simian |
|  | B-type -<br>SCN1B | not<br>applicable | yes | yes | yes | simian |

## STAR Methods

### EXPERIMENTAL MODEL AND SUBJECT DETAILS

#### Human tissue collection and preparation

The study on research involving human subjects was approved by the Ethics Committee Research of University Hospitals Leuven (UZ Leuven) (reference S61186). Prior to surgery, written informed consent was obtained.

Human cortical samples used for acute recordings and post-hoc staining as well as immunoprecipitations were resected from the temporal cortex during neurosurgery. All samples represented the lateral temporal neocortex and were obtained from patients who underwent amygdalohippocampectomy for medial temporal lobe seizures either due to hippocampal sclerosis, focal cortical dysplasia or low-grade mesial temporal tumours. They were obtained from patients, males and females, aged between 2 and 61 years old. Samples were collected at the time of surgery, immerged in ice-cold ACSF (NaCl 126 mM, NaHCO_3_ 26mM, D-glucose 10mM, MgSO_4_ 6mM, KCL 3mM, CaCl_2_ 1mM, NaH_2_PO4 1mM, 295-305mM, pH adjusted to 7.4, with 5% CO_2_/95% O_2_) and transferred immediately to the laboratory, with processing (slicing for electrophysiology or protein extraction) in an interval of 5-10 minutes. Slicing solution contained choline chloride 110mM, NaHCO_3_ 26mM, Na-ascorbate 11.6 mM, D-glucose 10mM, MgCl_2_ 7mM, Na-pyruvate 3.1 mM, KCl 2.5 mM, NaH_2_PO4 1.25mM, CaCl_2_ 0.5 mM; 300–315 mOsm, pH adjusted to 7.4, with 5% CO_2_/95% O_2_ and was ice-cold. Recovery solution was the same than the ACSF used for the transfer from hospital. Slicing was performed with a vibrating blade microtome or using a comprestome, and 300-µm slices were incubated for around 30 min at 32 °C in ACSF. Slices were then stored at around 20 °C until use for electrophysiological recordings.

Slices were immerged overnight in ice-cold PFA 4% (Histofix) and then stored in PBS azide 0.03% for post-hoc blind immunostaining.

Some human cortical samples used only for immunostaining were also obtained from UZ Leuven. These samples represented the lateral temporal, fronto-insular or temporo-parietal neocortex of patients, males and females, aged of 7, 11,14, 22 and 38 years old. They were immerged in the laboratory in sucrose 8% PFA 4% (pH = 7.4) during 3 hours and then conserved in PBS azide 0.03% before and after sectioning at the vibratome (80 µm thickness).

Other human cortical samples used only for immunostaining were originated from frozen specimen stored at the Netherlands Brain Bank (NBB), Netherlands Institute for Neuroscience, Amsterdam (reference 1256S). All material has been collected from donors for or from whom a written informed consent for a brain autopsy and the use of the material and clinical information for research purposes had been obtained by the NBB. They were obtained from control patients without known brain or cognitive disorders declared. These samples represented the superior and lateral occipital as well as superior parietal neocortex of individuals, males and females, aged of 24 and 45 years old. They were cut at the cryostat (25 µm thickness), stored at -80°C and post-fixated using sucrose 8% PFA 4% (pH = 7.4) during 30 minutes.

Other human cortical samples used only for immunostaining were originated from frozen specimen stored at the Edinburgh Brain Bank (reference BBN_2338) and King’s College London Brain Bank (references BBN_17052/17057/17060/17062), United Kingdom. All material has been collected from donors for or from whom a written informed consent for a brain autopsy and the use of the material and clinical information for research purposes had been obtained by the UK Brain Bank. They were obtained from control patients without brain or cognitive disorders declared. These samples represented the occipital and temporal neocortex of individuals, males and females, aged between 3 months, 4 months, 21 months and 16 years old. They were cut at the cryostat (25 µm thickness), stored at -80°C and post-fixated using sucrose 8% PFA 4% (pH = 7.4) during 30 minutes.

#### Non-human primate collection and preparation

Non-human primate samples used for immunostaining originated from frozen specimen stored at the Biomedical Primate Research Center, Rijswijk, The Netherlands. They were obtained from a 18 year-old male and a 17 year-old female chimpanzee and a 4-year old macaque rhesus. These samples represented the temporal neocortex. They were cut at the cryostat (25 µm thickness), stored at -80°C and post-fixated using sucrose 8% PFA 4% (pH = 7.4) during 30 minutes.

#### Animals

All mouse and rat experiments were performed with the approval of the KU Leuven Committee for animal welfare (protocol 2018/008, 089/2016 and 214/2017). Animals were housed under standard conditions (12 h light:12 h dark cycles) with food and water ad libitum. Mouse housing, breeding and experimental handling were performed according to the ethical guidelines of the Belgian Ministry of Agriculture in agreement with European community Laboratory Animal Care and Use Regulations (86/609/CEE, Journal officiel de l’Union européenne, L358, 18 December 1986). Embryos (aged E15.5) of the mouse strain ICR (CD1, Charles River Laboratory) were used for in utero electroporation. The plug date was defined as embryonic day (E)0.5, and the day of birth was defined as P0. Animals were processed between P24 and P32. The data obtained from all animals were pooled without discrimination of sexes for the analysis.

Data for this study are derived from a total of 112 mice of both sexes and 6 rats of both sexes.

#### Cell lines

HEK293TT human embryonic kidney cells were obtained from American Type Culture Collection (ATCC cat# CRL-11268). HEK293TT cells were grown in Dulbecco’s modified Eagle’s medium (DMEM; Invitrogen) supplemented with 10% fetal bovine serum (FBS; Invitrogen), 100mM Na-pyruvate, 8.9 mM NaHCO_3_, and penicillin/streptomycin (Invitrogen) and split using TrypLE™ Express Enzyme.

#### Protein and peptide reagents

FGF13A *E. coli* recombinant protein originates from Novus Biologicals (NBP2-35009). FGF13A ExonS and FGF13A ExonS-biotin (FGF13A amino acids 1-63) have been custom synthetized by ThermoFisher. LRRC37B-Fc recombinant protein was produced in Joris de Wit laboratory from HEK-293T cells. Serum extracted Fc protein was purchased at Jackson ImmunoReseach.

### METHOD DETAILS

#### Genome and transcriptome analysis

Encoding genes paralogs and orthologs originated from (Giannuzzi et al., 2012) and Ensembl gene trees (https://www.ensembl.org). Transcriptomic comparison between species is an analysis of data from Henrik Kaessmann laboratory (Heildelberg, Germany) described in (Cardoso-Moreira et al., 2020) and available at https://apps.kaessmannlab.org/evodevoapp/. Genes considered are the following: (macaque LRRC37-M7), ENSMUSG00000078632 (mouse LRRC37A) and ENSMUSG00000034239 (mouse GM884). Transcriptomic expression in the human cerebral cortex has been analysed from data available at the UCSC cell browser (autism data) taking control individuals available at https://cells.ucsc.edu/?ds=autism and published in (Velmeshev et al., 2019) and at M1 - 10X GENOMICS (2020) Allen Brain Single Cell Atlas (https://portal.brain-map.org/atlases-and-data/rnaseq) and published in (Bakken et al., 2021).

Copy number estimates for genes *LRRC37A*, *LRRC37A2*, *LRRC37A3*, and *LRRC37B* were obtained using QuicK-mer2 (Shen & Kidd, 2020) in windows of 500 unique *k*- mers. High-coverage whole-genome sequencing data form 2,504 unrelated individuals from five continental “super populations” (Byrska-Bishop et al., 2022) were downloaded in cram format and used as input for QuicK-mer2 with T2T-CHM13 (v1.0) as reference (Nurk et al., 2022). We genotyped overall gene CN as the mean CN across the gene body using a custom python script. CN-dotplots generated using the R package ggplot2.

#### Human and non-human primate cortex immunostaining

For immune fluorescent staining, human vibratome and cryosections as well as non-human primate sections were stained using Cy3 TSA amplification for LRRC37B. Briefly, slice were treated with tap water (5mn for vibratome sections, 1mn for cryosections), BLOXXAL reagent (3 hours for vibratome sections, 10mn for cryosections), three TNT washes (0.1 M TRIS-HCl, pH 7.5, 0.15 M NaCl, 0.3% Triton), TNB incubation for 2 hours and then incubation in TNB with rabbit anti-LRRC37B 1:1000 antibody which recognizes the LB specific domain (HPA015135, Merck) at 4°C (overnight for cryosections, 3 days for vibratome sections). After five washes with TNT, sections were incubated overnight at 4°C with anti-rabbit IgG antibody conjugated with HRP 1:100. Cy3 TSA reaction was performed after five washed with TNT (10mn reaction for vibratome sections, 3mn for cryosections). Slices were then transferred into the blocking solution (PBS 0.3% Triton, 5% horse serum, 3% BSA) and incubated for 1 hour. Brain slices were if required incubated at 4°C with mouse anti-ankyrin-G (1:500; MABN466, Merck) antibody (3 days for vibratome sections, overnight for cryosections). After three PBS washes, slices were incubated overnight at 4°C with donkey anti-mouse a488 or a647 and Hoechst (1:10000). For patched sections, brain slices were directly incubated overnight in PBS at 4°C containing streptavidin-a488 1:500 and Hoechst 1:10000. After three washes in PBS, brain sections were mounted on a slide glass with the mounting reagent (DAKO glycerol mounting medium) using #1.5 coverslips.

#### DNA constructs

LRRC37B cDNA originates from IRCMp5012D0514D (SourceBiosciences) whose sequence miss ExonS 4-5 which have been amplified by PCR from a cDNA library derived from GW18 fetal cortex (Suzuki et al., 2018) using the primers designed on the basis of the sequence of reference genome. The size of PCR fragment was confirmed and PCR fragment was subcloned into the BsmbI and EcorI restriction sites of the original cDNA by In Fusion cloning. LRRC37B cDNA has been inserted by PCR amplification and InFusion cloning into the multicloning site between CAG promotor and IRES in the lentiviral backbone pCIG (CAG-IRES-EGFP-WPRE, Addgene #122953) (Suzuki et al., 2018) and pCIG-LSL (CAG-LSL-IRES-EGFP-WPRE) described in (Iwata et al., 2020). Resulting pCIG-LRRC37B and pCIG-LSL-LRRC37B plasmids have been used compared to pCIG and p-CIG-LSL in in utero electroporation experiments.

LRRC37B cDNA has been PCR amplified adding a Cter HA tag and inserted into the pCIG backbone at the multicloning site between CAG promotor and IRES. Resulting pCIG-LRRC37B-HA plasmid has been used compared to pCIG in in utero electroporation experiments. All constructs were verified by DNA sequencing.

Predicted extracellular sequence lacking the signal peptide of LRRC37B (LRRC37B- ECTO) corresponding cDNA has been cloned by PCR amplification into a modified pCMV6-XL4 as described in (Apóstolo et al., 2020) leading to pLRRC37B-Fc using InFusion cloning. Fc-fusion protein contain a prolactin leader peptide (PLP) followed by an N-terminal FLAG tag, ectodomain of interest, a 3CPro cleavage site, and the dimeric human Fc domain. Similarly, LRRC37B-alkaline phosphatase (AP) fusion protein contain a leader peptide as described in (Apóstolo et al., 2020), a FLAG tag and ectodomain of interest. All constructs were verified by DNA sequencing.

Predicted extracellular sequence lacking the signal peptide of LRRC37B (LRRC37Becto) corresponding cDNA and truncated versions have been cloned by PCR amplification into pDisplay™ Mammalian Expression Vector (ThermoFisher V66020) using InFusion cloning resulting in pHA-LRRC37B-ECTO (amino acids 28 – 905), pHA-LRRC37B-ΔLRR-ECTO (amino acids 28 – 522: 748 – 905), pHA- LRRC37B-LRR-ECTO (amino acids 468 – 841), pHA-LRRC37BΔLB-ECTO (amino acids 186 - 905), pHA-LRRC37B-LB-ECTO (amino acids 28 – 520). All constructs were verified by DNA sequencing.

LRRC37A2 cDNA originates from OCABo5050B0130D (SourceBiosciences) and has been inserted into the pCIG backbone by PCR amplification adding a Cter HA tag or into the pCIG-LSL backbone by PCR amplification adding a Cter HIS tag, and InFusion cloning into the multicloning site between CAG promotor and IRES or LSL and IRES, respectively. All constructs were verified by DNA sequencing.

FGF13A cDNA plasmid originates from Origene (RC204164) and has been PCR amplified for insertion into pCIG plasmid by InFusion cloning. FGF13B, FGF13VY, FGF13core have been PCR amplified from FGF13A cDNA with primers targeting the core domain of FGF13 and with ExonS B or VY in the 5’ primer. All constructs were verified by DNA sequencing.

pCMV-macaqueLRRC37B-HA (ORF XP_028692824.1) and pCMV-chimpanzeeLRRC37B-HA (ORF CK820_G0028539) have been synthetized by GenScript and inserted in pcDNA3.1(+)-C-HA backbone (Addgene #128034). All constructs were verified by DNA sequencing.

pCMV-SCN1B-FLAG, plasmid originates from Origene (RC209565, RC215868, RC219002). Predicted extracellular sequence lacking the signal peptide of SCN1B (SCN1Becto) corresponding cDNA has been cloned by PCR amplification into pDisplay™ Mammalian Expression Vector (ThermoFisher V66020) using InFusion cloning resulting in pHA-SCN1Becto. All constructs were verified by DNA sequencing.

pCMV-SCN8A-IRES-Scarlet plasmid originates from Addgene (#162280) and has been described in (DeKeyser et al., 2021). pCAG-cre is a gift from Franck Polleux laboratory (United States) described in (Hand et al., 2005). pCAG-GEPH.FingR- tdTomato-IL2RGTC is a gift from Juan Burrone (United Kingdom), derived from pCAG_GPHN.FingR-mKate2-IL2RGTC (Addgene #46297) described in (Gross et al., 2013). pGPR158_ECTO, pSLIRTK2_ECTO and pLRRTM1_ECTO were previously described (Condomitti et al., 2018; Schroeder et al., 2018). pCD4_ECTO is a gift from Luís Ribeiro (Joris de Wit’s laboratory), with the cDNA of the predicted extracellular domain of CD4, originating from pCMV-CD4 (Addgene #51604) described in (Raissi et al., 2013), inserted into the pDisplay™ Mammalian Expression Vector (ThermoFisher V66020).

#### Live staining of HEK-293T cells

HEK-293T cells were split on coverslips and have been transfected with 500ng of each construct total amount (1ug) of DNA (pCIG + pCIG-humanLRRC37B-HA or + pCMV- chimpanzeeLRRC37B-HA or + pCMV-macaqueLRRC37B-HA or + pCIG- humanLRRC37A2-HA), using XtremeGene9 transfection reagent.

For live staining, 48 hours after transfection, medium was washed with cold PBS and then rabbit anti-LRRC37B (1:1000, as described above) was applied in PBS for 1h at 4°C. This was followed by washes in PBS, fixation 30mn with PBS PFA 4% at 4°C. After washes in PBS, cells were blocked for 1h in PBS 5% HS 3% BSA, and then with 1:1000 donkey anti-rabbit Cy3 and Hoechst 1:10000 for 2 hours in the same solution at room temperature.

For staining after fixation and permeabilization, cells were washed with cold PBS and fixed 30mn with PBS PFA 4% at 4°C. After washes in PBS, cells were blocked for 1h in PBS 5% HS 3% BSA, and then with rabbit anti-HA (1:1000; CST 3724S, Bioke) overnight at 4°C in the same solution. After washes in PBS, 1:1000 donkey anti-rabbit Cy3 and Hoechst was applied for 2 hours in the same blocking solution at room temperature.

Coverslips were mounted on a slide glass with the mounting reagent (DAKO glycerol mounting medium).

#### In utero electroporation

Barrel cortex of E15.5-day-old embryos of timed-pregnant CD1 mice were unilaterally electroporated with lentiviral plasmids. Briefly, the dam was anesthetized with isoflurane following buprenorphine and carprofen injection, and the uterus exposed. A solution of 1-2 μg/μl DNA and 0.01% fast green dye was injected into the embryonic lateral ventricle with a heat-pulled glass capillary. For immunostainings and image analysis, pCIG-LSL plasmids (1000 ng/µL) were used together with pCAG-cre (15 ng/ µL) eventually with pCAG-GEPH.FingR-tdTomato-IL2RGTC while for electrophysiology pCIG plasmids were used (1 μg/μl) (see above). The embryo’s head was then placed between the paddles of pair of tweezer electrodes with the cathode lateral to the filled ventricle and five 50 ms, 30 V pulses were delivered with an interval of 950ms by a BTX830 electroporator (Harvard Apparatus). After electroporation, the uterus was replaced, the incision sutured closed and placed on a heating plate until recovery.

#### Mouse cortex processing and immunostaining

Mouse P28 animals were perfused transcardiacally with ice-cold sucrose 8% PFA 4%. Brains were dissected and soaked in the same fixative for 3 hours, then stored in PBS azide. Then they either have been sectioned in 80 μm thickness using vibratome or 50 μm thickness using cryostat after dehydration in sucrose 30% and freezing in OCT. Slices were transferred into the blocking solution (PBS 0.3% Triton, 5% horse serum, 3% BSA) and incubated for 1 hour. Brain floating slices were incubated 3 days at 4°C with primary antibodies: rabbit anti-LRRC37B (1:1000, as described above), mouse anti-ankyrin-G (1:500, as described above), mouse anti-FGF13A (1:500; MA5-27705, ThermoFisher), chicken anti-EGFP (1:1000; ab13970, Abcam), rat anti-mCherry which recognizes tdTomato (1:1000; M11217, ThermoFisher), mouse anti-pan-NAVα (1:500; S8809, Merck). For VGAT staining (guinea pig anti-VGAt 1:500; 131 004, Synaptic Systems), stainings have been done sequentially in blocking solution PBS 1% Triton, 5% horse serum, 3% BSA. After three PBS washes, slices were incubated overnight at 4°C with secondary antibodies in PBS: donkey anti-rabbit Cy3, anti-rabbit a594, anti-mouse a647, anti-chicken a488, anti-rat Cy3, anti-guinea pig a647 (1:1000 or 1:250 for cryosections used for STED imaging) and Hoechst (1:10000). After three washes in PBS, brain sections were mounted on a slide glass with the mounting reagent (DAKO glycerol mounting medium) using #1.5 coverslips.

#### Image acquisition

Confocal images were obtained with Zeiss LSM880 and LSM900 driven by Zen Black and Blue softwares equipped with objectives 10x, 20x, oil immersion 25x and oil immersion 40x, AiryScan system and argon, helium-neon and 405 nm diode lasers.

STED single focal section images were obtained with an Abberior system with Olympus IX83 body equipped with 100x oil immersion, 480, 532, 640 nm excitation lasers and 595nm 775 nm depletion lasers. STED pictures were deconvoluted using Huygens deconvolution software. Except if specified, representative pictures are maximum projections.

#### Image analysis

AIS intensity profile was done on maximum projection pictures in Matlab as described in (Grubb and Burrone, 2010). In mouse, the beginning of the AIS was set using the EGFP channel (starting from the soma). In human, ankyrin-G was used to set the beginning of the axon.

Puncta and area quantification have been done using Fiji. On average, 10 focal sections (0.4 µm thickness) maximum projection was used for quantification. For STED imaging, one focal section was used only with on average 10µm length in the proximal part of the AIS. EGFP was used to delineated the soma and the AIS (30µm starting from the soma). Binarized pictures of the gephyrin-tdTomato, VGAT, FGF13A have been used to quantify manually the number of puncta or their positive area.

#### Electrophysiological recordings

For mouse experiments, coronal slices were prepared from postnatal day P24–32 animals. Briefly, after decapitation, the brain was quickly removed and transferred into ice-cold cutting solution (in mM): 87 NaCl, 2.5 KCl, 1.25 NaH_2_PO_4_, 10 glucose, 25 NaHCO_3_, 0.5 CaCl_2_, 7 MgCl_2_, 75 sucrose, 1 kynurenic acid, 5 ascorbic acid, 3 pyruvic acid, pH 7.4 with 5% CO_2_/95% O_2_, and whole brain coronal slices (250 µm) were cut using a vibratome (VT1200, Leica Biosystems). Afterward, slices were transferred to 32 °C cutting solution for 45 min to recover and finally maintained at room temperature until used for recordings.

Human cortical samples were collected at the time of surgery, immerged in ice-cold ACSF (NaCl 126 mM, NaHCO3 26mM, D-glucose 10mM, MgSO4 6mM, KCL 3mM, CaCl2 1mM, NaH2PO4 1mM, 295-305mM, pH adjusted to 7.4, with 5% CO2/95% O2) and transferred immediately into the laboratory, with processing (slicing for electrophysiology or protein extraction) in an interval of 5-10 minutes. Slicing solution contained choline chloride 110mM, NaHCO3 26mM, Na-ascorbate 11.6 mM, D-glucose 10mM, MgCl2 7mM, Na-pyruvate 3.1 mM, KCl 2.5 mM, NaH2PO4 1.25mM, CaCl2 0.5 mM; 300–315 mOsm, pH adjusted to 7.4, with 5% CO2/95% O2 and was ice-cold. Recovery solution was the same than the ACSF used for the transfer from hospital. Slicing was performed with a vibrating blade microtome or using a comprestome, and 300-µm slices were incubated for around 30 min at 32 °C in ACSF. Slices were then stored at around 20 °C until use for electrophysiological recordings.

For recordings, mouse and human brain slices were continuously perfused (32-34°C) in a submerged chamber (Warner Instruments) at a rate of 3–4 ml/min with (in mM): 127 NaCl, 2.5 KCl, 1.25 NaH_2_PO_4_, 25 NaHCO_3_, 1 MgCl_2_, 2 CaCl_2_, 25 glucose at pH 7.4 with 5% CO_2_/95% O_2_. Whole-cell patch-clamp recordings were done using borosilicate glass recording pipettes (resistance 3.5–5 MΩ, Sutter P-1000), using a double EPC-10 amplifier under control of Patchmaster v2 x 32 software (HEKA Elektronik, Lambrecht/Pfalz, Germany). The following internal medium was used (in mM): 135 K-Gluconate, 4 KCl, 10 HEPES, 4 Mg-ATP, 0.3 Na_2_-GTP, 10 Na_2_- phospocreatine, 3 biocytin (pH 7.3). Cell intrinsic properties were recorded in current clamp, while sEPSCs and sIPSCs were recorded in voltage clamp at -70 mV and 0 mV, respectively. Currents were recorded at 20 Hz and low-pass filtered at 3 kHz when stored. The series resistance was compensated to 75-85%. Cell intrinsic properties were analyzed using Fitmaster (HEKA Elektronik, Lambrecht/Pfalz, Germany), spontaneous input was analyzed using Mini Analysis program (Synaptosoft). For protein/peptide bath application experiments we used consecutive repeats of recordings (control followed by recordings 3 to 16 minutes after application), 10 minutes application is plotted.

For initial intracellular application experiments, 50 nM FGF13A were added to the intracellular pipette solution and standard approach procedures were used. Next, to minimize FGF13A exposure of the slice, a fluorescent dye was added to the intracellular medium (Alexa568, 5 µM) to visually regulate minimal outflow of intracellular solution from the pipette during the approach and cell-attached phase of recordings. In both experiments (no application versus application), recordings were started 3 minutes after establishing whole-cell configuration, to allow infusion of FGF13A into the cell.

Phase plot analysis compares the rate of change (first derivative) in voltage during APs (y-axis) to the membrane voltage (x-axis). The first current step (1 sec) to initiate AP(s) was used for phase plot analysis. The AIS, somatic and repolarization voltages were determined as the absolute membrane potential measured at the respective peaks.

#### Elisa assay

An ELISA-based assay (Ozgul et al., 2019) was used to identify the interaction between ectodomain (as defined in the Uniprot database) of 920 cell surface or secreted proteins cloned in frame with an Fc domain against LRRC37B-ECTO-AP fusion as described in (Apóstolo et al., 2020). Horseradish peroxidase (HRP) conjugated anti-Fc antibody develops a blue colour if the prey (FGF13A) remain bound to the bait (LRRC37B) after the washes. After the initial identification, the experiment has been repeated 3 times using triplicate wells. The library contained AIS proteins or proteins coded by genes enriched in chandelier interneurons as described in (Bakken et al., 2021; Favuzzi et al., 2019; Leterrier, 2016): *ALCAM, CDH4, CDH6, CDH11, CNTNAP5, DPP10, FGF13 isoform 1 (FGF13A), FSTL5, ITGAV, ITGA6, LRRN1, LRRN2, NFASC, OLFM3, PCDH19, PCSK2, ROBO1, SGCD, SLITRK1, SLITRK5, TENM4, THSD7A, UNC5B*. They were all negative except FGF13A.

#### Protein extraction and immunoprecipitations

HEK293T have been transfected with 500-2000ng of DNA using XtremeGene9 transfection reagent. 72 hours after transfection, cells were lysed in lysis buffer (50mM Tris-HCl, pH 7.5, 1% sodium deoxycholate, 0.1% Triton, proteinase inhibitors) on a wheel for one hour in a cold room. When required, FGF13A and its ExonS where applied in the culture medium 5 hours before protein extraction.

EGFP positive area of P17 mouse cortex has been dissected using forceps in cold PBS of brains after cervical dislocation. Cortices have been homogenized in homogenization buffer (50mM Tris-HCl, pH 7.5, 1% sodium deoxycholate, proteinase inhibitor) using a Dounce homogenizer. After Triton addition (0.1% final concentration), samples have been rotated on a wheel for one hour in a cold room.

Human cortices (from 14yo-48yo patients) have been homogenized in homogenization buffer (50mM Tris-HCl, pH 7.5, 1% sodium deoxycholate, proteinase inhibitor) using a Dounce homogenizer. After Triton addition (0.1% final concentration), samples have been rotated on a wheel for one hour in a cold room.

After incubation, samples were centrifugated 25mn 16000g and lysates transferred in a new tube with addition of NaCl (final concentration 150 mM). Samples were incubated overnight on a wheel in a cold room with HA magnetic beads or protein A magnetic beads coupled with 1ug of mouse anti-pan-NAVα (as described above) or rabbit anti-FLAG (ab1162, Abcam) or mouse anti-FGF13A (as described above) or mouse IgG (ThermoFisher) antibodies. Beads were washed 4 times with the washing solution (50mM Tris-HCl, pH 7.5, 1% sodium deoxycholate, 0.1% Triton,150 mM) and one time with PBS. Subsequently, samples were eluted in 2x Laemmli buffer at 95°C. Medium was centrifugated 25mn 16000g and diluted to final 1x Laemmli buffer at 95°C. The input (in 1x Laemmli buffer), medium, and immunoprecipitated samples were run in NUPAGE 12% Bis-Tris Protein Gel at the voltage of 90V for 2 hours in MOPS buffer and then transferred to PVDF Blotting Membrane at the voltage of 100V for 100 minutes. The membrane was blocked in the buffer (5% skim milk and 0.1% Tween20 in TBS) for 1 hour at room temperature and subsequently incubated in the blocking buffer containing rabbit anti-HA (1:1000, as described above), rabbit anti- LRRC37B (1:1000, as described above), mouse anti-FGF13A (1:1000, as described above), mouse anti-pan-FGF13 (1:1000; PA5-27302, ThermoFisher), mouse anti-beta-actin (1:5000; MA1-140, ThermoFisher), mouse anti-pan-NAVα (1:500, as described above), rabbit anti-FLAG (1:1000; as described above), rabbit anti-SCN1B (1:500; CST 13950S, Bioké) antibodies overnight at 4°C, followed by the incubation in the blocking solution containing secondary antibody anti-Rabbit or Mouse IgG antibody conjugated with HRP at room temperature for 1 hour. Pierce ECL Western Blotting Substrate was used for signal detection.

#### Binding assay & affinity approximation

To estimate the FGF13A – LRRC37B affinity on the cell surface, a binding assay approach has been used as described in (Savas et al., 2015). Briefly, HEK-293T cells were plated on 10cm plates, non-transfected (6 times) or transfected with the pdisplay empty vector (6 times), LRRC37B_ECTO (6 times), LRR_ECTO (3 times),

ΔLRR_ECTO (3 times)(described above), cultured for 24 hours, gently trypsinized and re-plated on 24 well plates and cultured for an additional 24 hours. Live cells were incubated with FGF13A ExonS-biotin at 0, 1, 5, 10, 50, 100 nM, fixed, probed with a streptavidin-HRP and reacted with TMB. The reaction was stopped with 1N HCl and transferred to 96 well plates and the absorbance was measured on a plate reader at 450nm. All saturation binding calculations were performed with GraphPad Prism, One site –Specific binding, non-linear fit curve.

#### Pull down from rat brain extracts

Fc-Protein Purification for Mass-spectrometry was performed as described previously (Savas et al., 2014). LRRC37B (aa 28-905, containing the entire ectodomain) ecto-FC protein was produced by transient transfection of HEK293T cells using PEI (Polysciences). Six hours after transfection, media was changed to OptiMEM (Invitrogen) and harvested 5 days later. Conditioned media was centrifuged, sterile-filtered and run over a fast-flow Protein-G agarose (Thermo-Fisher) column. After extensive washing with wash buffer (50 mM HEPES pH 7.4, 300 mM NaCl and protease inhibitors), the column was eluted with Pierce elution buffer. Eluted fractions containing proteins were pooled and dialyzed with PBS using a Slide-A-Lyzer (Pierce) and concentrated using Amicon Ultra centrifugal units (Millipore). The integrity and purity of the purified ecto-Fc proteins was confirmed with SDS-PAGE and Coomassie staining, and concentration was determined using a Bradford protein assay.

Affinity chromatography experiments were performed as previously described (Savas et al., 2014). Crude synaptosome extracts were prepared from 2-3 P21-22 rat brains per condition, homogenized in homogenization buffer (4 mM HEPES pH 7.4, 0.32 M sucrose and protease inhibitors) using a Dounce homogenizer. Homogenate was spun at 1,000 x g for 10 minutes at 4°C. Supernatant was spun at 14,000 x g for 20 minutes at 4°C. P2 crude synaptosomes were re-suspended in Extraction Buffer (50mmHEPES pH 7.4, 0.1MNaCl, 2 mM CaCl2, 2.5 mM MgCl2 and protease inhibitors), extracted with 1% Triton X-100 for 2 hours and centrifuged at 100,000 x g for 1 hour at 4 to pellet insoluble material. Fast-flow Protein-A Sepharose beads (GE Healthcare) (250 µl slurry) pre-bound in Extraction Buffer to 100 µg human Fc or LRRC37-Fc were added to the supernatant and rotated overnight at 4°C. Beads were packed into Poly-prep chromatography columns (BioRad) and washed with 50 mL of high-salt wash buffer (50 mM HEPES pH 7.4, 300 mM NaCl, 0.1 mM CaCl2, 5% glycerol and protease inhibitors), followed by a wash with 10 mL low-salt wash buffer (50 mM HEPES pH 7.4, 150 mM NaCl, 0.1 mM CaCl2, 5% glycerol and protease inhibitors). Bound proteins were eluted from the beads by incubation with Pierce elution buffer and TCA-precipitated overnight. The precipitate was re-suspended in 8 M Urea with ProteaseMax (Promega) per the manufacturer’s instruction. The samples were subsequently reduced by 20-minute incubation with 5mM TCEP0 (tris(2carboxyethyl)phosphine) at RT and alkylated in the dark by treatment with 10 mM Iodoacetamide for 20 additional minutes. The proteins were digested overnight at 37°C with Sequencing Grade Modified Trypsin (Promega) and the reaction was stopped by acidification. Mass spectrometry analysis was performed by the VIB Proteomics Core (Ghent, Belgium).

#### Statistical analysis

Data are represented as mean + standard error of the mean (SEM), individual values with mean, or if when detailed in the Figures with median + SE median. Unpaired Mann-Whitney test or paired Wilcoxon tests have been used for assessing the significance of differences in the analyses containing two conditions and two-way ANOVA in the analyses containing more than three conditions using GraphPad. See each figure for details.

For mouse image quantification data, we used at least 3 neurons per animal, 3 animals per litter (littermates comparison) and at least 3 litters. For mouse IUE electrophysiological data we used 9-15 animals per group from four litters. Peptide application was done on 2-7 animals per condition.

For co-immunoprecipitations, experiments with HEK-293T cells have been performed >3 times, experiments from mouse *in vivo* samples > 2 times and human *in vivo* samples >3 times.

For human AIS intensity profile data we analysed 60 neurons per patient, 3 patients (7-38 years old); and for electrophysiology, 4-6 neurons per patient and 7 patients (4- 62 years old).

